# Widespread male sterility and trioecy in androdioecious *Mercurialis annua*: an account of its distribution, its genetic basis, and estimates of its morph-specific fitness components

**DOI:** 10.1101/2024.07.31.605989

**Authors:** M. T. Nguyen, T. Martignier, J. R. Pannell

## Abstract

**Premise:** Angiosperms range from hermaphroditism through gynodioecy and androdioecy to dioecy. ‘Trioecy’, where females and males coexist with hermaphrodites, is rare. Recently, trioecy was documented in hexaploid populations of the wind-pollinated herb Mercurialis annua in Spain.

**Methods:** We surveyed the frequency of males, hermaphrodites and females in *M. annua* across its distribution in the Iberian Peninsula, tracked sex-ratio variation in several populations over consecutive generations, and assessed evidence for pollen limitation. In a common garden, we estimated male, female and hermaphrodite fitness. We used controlled crosses to infer the genetic basis of male sterility. Finally, we compared predictions of a deterministic model with the distribution of observed sex ratios in the field based on our fitness estimates and the inferred genetics of sex determination.

**Key results:** Trioecy is widespread in Spanish and Portuguese populations of *M. annua*. Males are determined by a dominant (Y-linked) allele, and female expression results from the interaction between cytoplasmic male sterility and multiple nuclear male sterility restorers partially linked to the male determiner. Male pollen production is approximately 12 times while female seed production is less than 1.12 times that of hermaphrodites. The distribution of sex ratios in natural populations conforms with predictions of our deterministic simulations.

**Conclusions:** Our study documents and accounts for a clear case of trioecy in which sex is determined by both maternally and biparentally inherited genes.

## INTRODUCTION

Sexual systems vary substantially among flowering plants. In patterns of allocation to sex, they range from hermaphroditism and monoecy, by far the most common sexual systems in which all individuals allocate to both male and female functions, either within each flower (‘hermaphroditism’) or in separate flowers (monoecy), to dioecy, in which individuals are either male or female (Barrett, 2002; Pannell and Jordan, 2022). While dioecy is much rarer than hermaphroditism, accounting for approximately 6% of angiosperms (Renner and Ricklefs, 1995), it has evolved on numerous occasions independently and occurs in about half of all angiosperm families. Gynodioecy and androdioecy, in which females or males, respectively, co-occur with hermaphrodites, lie between these two sex-allocation extremes and are rarer (Lewis, 1941; Lloyd, 1975; Charlesworth and Charlesworth, 1978), especially androdioecy, which has been found in only a handful of species (Charlesworth, 1984; Pannell, 2002). Trioecy, in which males and females both co-occur with hermaphrodites, is also very rare (Yampolsky and Yampolsky, 1922; Godin, 2022). Although trioecy is rare, an understanding of its biology may lead to important insights relevant to the evolution of sexual systems (Charlesworth and Charlesworth, 1978; Maurice et al., 1994; Schultz, 1994) and sex allocation (Pannell and Jordan, 2022) more generally, as well as to the genetic architecture of sex determination (Takahashi et al., 2023).

The genetic basis of the determination of males and females, and the evolution and maintenance of androdioecy and gynodioecy from hermaphroditism, have been well-investigated theoretically and empirically (Charlesworth and Charlesworth, 1978; Pannell and Jordan, 2022). Males are predominantly determined by a Y chromosome in the nucleus (Weestergaar, 1958; Charlesworth 1984, 2002), and need to produce at least twice the amount of pollen as hermaphrodites for androdioecy to be maintained (Charlesworth and Charlesworth, 1978). Females may be determined by nuclear genes, but most gynodioecious species have females determined by cytoplasmic genes (Kaul, 1988). Nuclear-determined females need to produce at least twice the number of seeds as hermaphrodites to establish gynodioecy in outcrossing populations (Lewis, 1941; Charnov et al., 1976, Charlesworth and Charlesworth, 1978), whereas cytoplasmic-determined females only need to produce more seeds than hermaphrodites for gynodioecy to evolve (Lewis, 1941; Lloyd 1975). In both cases, females may invade hermaphroditic populations under less stringent conditions if hermaphrodites are partially selfing and selfed progeny suffer from inbreeding depression (Charlesworth and Charlesworth, 1978). While we have a good understanding of the genetic basis and the conditions under which hermaphroditism, gynodioecy, and androdioecy can be maintained by natural selection, we have a much poorer understanding of the possible maintenance of trioecy, both from a theoretical point of view, as well as in terms of empirical observations consistent with the existence of three different sex-allocation strategies in wild populations.

Theoretical modeling has largely failed to identify conditions for the evolution and maintenance of trioecy. In a model assuming biparental inheritance of genes affecting sex allocation, Charnov et al. (1976) concluded that trioecy could not be maintained in a stable equilibrium. Gregorius et al. (1983) proposed a one-locus biallelic model and found that trioecy could be maintained only if males and females were determined by recessive sterility alleles. Wolf and Takebayashi (2004) considered a model for trioecy with three alleles segregating at a single locus and further introduced the possibility of pollen limitation, yet they too concluded that trioecy could not be maintained. In contrast, in the only model to date that has found conditions for evolutionary stable trioecy, Maurice and Fleming (1995) found trioecy could be maintained as long as pollen limitation reduced the fitness of females more than that of hermaphrodites.

Given the theoretical predictions that trioecy should only be maintained under restricted conditions, we ought to be cautious in evaluating claims of its occurrence in nature. Ultimately, trioecy ought only to refer to populations in which males, females and hermaphrodites represent discrete sexual strategies (Sakai and Weller, 1999). However, for most populations that have been labeled trioecious in the literature, we lack sufficient data to evaluate how discrete the putative three phenotypes are. Many cases should probably more properly be described as subdioecious or dioecious, with males and/or females displaying inconstant or ‘leaky’ sex expression by producing a few flowers of the opposite sex (Lloyd, 1980; Ehlers and Bataillon, 2007). Vitale and Freeman (1986) referred to *Spinacea oleraceae* as trioecious, but the hermaphrodites are likely just leaky males and males, as the authors also noted that the sex expression is labile. The complex sex expression observed in the tree *Fraxinus excelsior*, in which plants have been observed to produce either only male flowers, only female flowers or bisexual flowers, has been labelled trioecious (Albert et al., 2013), but it is now known that the species displays a type of subdioecy associated with an unusual diallelic self-incompatibility system (Saumitou-Laprade et al., 2018). Trioecy has also been claimed for the shrub *Coccoloba cereifera*, but the case may rather be an example of the coexistence of sexual with asexual reproduction, with the seed production of females being apparently agamospermous (Silva et al., 2008). The usually dioecious willow *Salix myrsinifolia* includes some populations with bisexual catkins that have been labeled trioecious, but no data on the distribution of gender among individuals in these populations have been published, and this example, too, might be another interesting case of subdioecy with inconstant sex expression, perhaps enhanced by selection for reproductive assurance during colonization phases, as suggested by the authors (Mirski and Brzosko, 2015). The aquatic herb *Sagittaria latifolia* has some populations with males, females and monoecious individuals (Sarkissian et al., 2001), but variation in sex expression in these populations tends to be continuous rather than discrete, and they may be the result of gene flow between dioecious and monoecious populations that are well characterized in the species complex (Yakimowski and Barrett, 2014). While many reported cases of trioecy are thus probably interesting examples of subdioecy or other systems with complex sex expression, a few species may, in fact, be characterized by trimodal variation in sex allocation and may thus qualify as meaningfully trioecious. One prime example of trioecy is the cactus species *Pachycereus pringlei* (Fleming et al., 1994). The female flowers of *P. pringlei* are smaller than hermaphrodites, with smaller, darker, pollenless anthers, while male flowers have fully developed stigmas and styles, but their ovaries lack ovules or contain few ovules developed into sterile fruits (Fleming et al., 1994). Similarly, the trioecious cactus species *Opuntia robusta* also has three distinct sex-morphs: females, with short styles and white, empty anthers; males, with long styles, atrophied stigmas, and reduced ovaries containing no or few atrophied ovules; and hermaphrodites, with intermediate styles and anthers and ovaries similar to those of males and females, respectively (Del Castillo and González-Espinosa, 1988; Del Castillo and Argueta, 2009).

An additional case of trioecy was described and studied by Perry *et al*. (2012) for several populations of the wind-pollinated annual herb *Mercurialis annua* in south-eastern Spain. In the Iberian Peninsula more generally, populations of *M. annua* vary between monoecy and androdioecy, where males co-occur with monoecious hermaphrodites (Pannell, 1997a, c, b; Pannell et al., 2008). For simplicity, we often refer to these individuals as ‘hermaphrodites’, as they transmit genes to progeny via both seeds and pollen. In the populations studied by Perry *et al*. (2012), males co-occurred with both fully fertile hermaphrodites as well as with ‘hermaphrodites’ expressing male sterility (i.e., functional females), resulting in a functional trioecious sexual system with females that differ discretely from hermaphrodites. The females in trioecious populations of *M. annua* produce somewhat smaller male flowers without anthers (or with small, sterile anthers) and are in all other ways phenotypically similar to fertile hermaphrodites. In contrast with the finding that sex ratios (male frequencies) in androdioecious *M. annua* depended on the density of plants in the previous (parental) generation (Dorken and Pannell, 2008), Perry *et al*. (2012) found no evidence for such density dependence in the trioecious populations of *M. annua*. This finding was surprising because the selfing rate is density-dependent in androdioecious *M. annua* (Eppley and Pannell, 2007a; Dorken and Pannell, 2008; Korbecka et al., 2011), is known to affect the sex ratio (Lloyd, 1975; Pannell, 1997a), and might be expected to do so under trioecy, too. Interestingly, seed set by females in dioecious populations of *M. annua* is sensitive to density, so trioecious populations might satisfy conditions for the maintenance of hermaphrodites with males and females, as predicted by Maurice and Fleming (1995). However, there is yet no evidence for pollen-limited seed production in trioecious *M. annua*, and Perry *et al*. (2012) could offer no satisfactory explanation for the insensitivity of trioecious sex ratios to density.

In the present article, we provide a more comprehensive assessment of the case of trioecy in *M. annua* in the Iberian Peninsula, addressing not only its geographic distribution and sex ratio, but also key components of reproductive success of the three sex-morphs and the genetic basis of morph sex determination. Since Perry *et al*.’s (2012) study, we have identified male sterility in many other populations of otherwise monoecious or androdioecious *M. annua*, principally in southern Spain, but in fact also widely around the Peninsula. Here, we first assess the distribution and frequency of males, hermaphrodites and females (i.e., male sterility) in *M. annua* across the distribution of the species’ range in the Iberian Peninsula on the basis of a large-scale survey. We also track sex-ratio variation in several natural trioecious populations in southern Spain over consecutive generations and assess evidence for pollen limitation as a function of density in a controlled experiment. In a common garden experiment, we estimate fitness components for males, females and hermaphrodites. Finally, we use controlled crosses to infer the genetic basis of male sterility in trioecious *M. annua*. We integrate our estimates of fitness components and the inferred genetic architecture of sex determination into deterministic calculations to compare the distribution of observed sex ratios in the field with those predicted by theory. Taken together, our results provide a detailed view of the maintenance of male sterility in androdioecious *M. annua* and the most comprehensive case study of trioecy in flowering plants to date.

## MATERIALS AND METHODS

### Study species —

*M. annua* (Euphorbiaceae) is an annual herb that grows in disturbed habitats around the Mediterranean Basin, central and western Europe (Durand, 1963). Although the species is dioecious (and diploid) in northern Iberia and throughout most of Europe, its polyploid hexaploid populations are variously monoecious or androdioecious throughout most of coastal Spain and Portugal (Pannell, 1997c; Pannell et al., 2014). Recently, Perry et al. (2012) described hexaploid populations in southeastern Spain in which individuals that superficially appeared monoecious were male-sterile (i.e., female), occurring at low frequencies (< 40%). In the current study, we describe and analyze the distribution of male sterility in *M. annua* more broadly across the Iberian Peninsula.

### Spatial distribution of male sterility in M. annua —

To characterize the distribution of male sterility in hexaploid *M. annua*, we surveyed populations of the species over two seasons (2015 and 2018) along a transect from Valencia in the southeast, clockwise around the Iberian Peninsula, to Santiago de Compostela in northwestern Spain. For most of the survey, we looked intensively for populations along roadsides at predetermined sites every 50 km. Between Gibraltar and Motril in southern Spain, where male sterility was observed to be more frequent, we surveyed populations at shorter spatial intervals (at least 500 m between adjacent populations). In each population, we estimated its size in terms of the total number of individuals, its surface area (and thus its density), and the numbers of fertile and sterile males and male-fertile and male-sterile monoecious individuals in a sample of at least 100 individuals; in what follows, we refer to the former two phenotypes as ‘males’ and ‘sterile males’, and to the latter two phenotypes as ‘hermaphrodites’ and ‘females’.

We determined whether male sterility is more likely in androdioecious than monoecious populations using a Wilcoxon test, defining androdioecy as a population with males at a frequency of > 0.025. Further, we determined whether the frequency of male sterility depends on plant density and male frequency on the basis of a generalized linear model with a binomial error distribution.

### Variation in the frequency of sexual phenotypes among seasons —

To characterize the extent to which the frequencies of sexual phenotypes might fluctuate from one year to the next, we established permanent plots at eight sites in southern Spain, between Gibraltar and Motril, and recorded demographic and phenotype variables in March each year over three consecutive seasons (2016, 2017 and 2018; Table S2). We divided the populations into marked quadrats of 1 m^2^. In each square, we recorded the number of plants and the frequency of each sexual phenotype. Out of the eight populations investigated in 2016, two were extinct in 2017, and a further population had become extinct in 2018. For the populations that had increased in size, we also scored plants in additional quadrats over the expanded area.

### Expression of sexual phenotypes in a common garden —

We investigated the differences in components of fitness between males, females and hermaphrodites in a common garden in the summer of 2016, using seeds from a population sampled near Motril (lat. 36.72791; long. −3.53112). The seeds were germinated in seed trays in a peat-based substrate. A month after germination, when the plants had begun to flower, they were transplanted into 1.3 L pots in a substrate comprising 25% topsoil, 10% vegetable fiber, 45% mineral component, and 20% compost. In each pot, we transplanted one male, one female, and one hermaphrodite. The pots were separated into three experimental blocks of 70 pots each (210 plants per block) and placed outside. Plants were harvested after two months of growth in their pots. We recorded plant height, above-ground biomass, the biomass of male flowers per plant, and the number and biomass of seeds per plant. Above-ground plant biomass and male-flower biomass were measured after a drying period of 48h at 60°C. We used a linear mixed model to test for differences among the sexes, with plant sex as a fixed factor and pot, block and female parent as random factors, and pot nested in block.

### Estimation of pollen limitation —

To determine the extent to which plants growing at low density may be pollen-limited in their seed set, we established females or hermaphrodites in circles around a single pollen source of males growing in the center and counted the numbers of developed fruits and undeveloped fruits (inferred to have been pollinated and unpollinated, respectively). Plants for this experiment were grown from seeds sampled from a population near Motril in southern Spain (lat. 36.72791; long. −3.53112), using plant culture protocols, as described above. One week after transplantation, the plants were separated into three replicates and placed outside. Each replicate consisted of one central group and two rings of plants. The central group was composed of four males, four females and four hermaphrodites (12 plants in total), randomly placed as closely as possible. The first ring of plants was composed of four females and four hermaphrodites that were placed 0.5 m away from the closest plant in the center and 0.5 m away from the closest plant in the same ring. The second ring of plants was placed similarly, but at 1.5 m from the center and with 1.5 m between plants on the same ring. On each ring, females and hermaphrodites were alternated. Thus, each replicate was composed of 28 plants, and the replicates were separated by 10 meters. The plants were left to grow for three months and were harvested in mid-autumn to estimate levels of pollen limitation. We estimated pollen limitation as the number of female flowers not developing fruits divided by the total number of female flowers on an individual. We also measured the biomass of male flowers for each plant.

To compare phenotypic traits (male flower weight per plant biomass, seed number and seed biomass per plant biomass) among sexes (male, female and hermaphrodite), we constructed linear mixed models (LMM), with plant sex as a fixed factor and replicate plot as a random factor. To analyze whether pollen limitation and seed production differed among sexes and with distance from the central male pollen source, we used LMM with distance from the center, sex and their interaction as fixed effects and replicate plot as a random effect. We then simplified our model by dropping first the interaction between the distance from the center and sex, based on AIC criterion, and were left with an LMM model with two fixed effects (sex and distance) and one random effect (plot replicate).

All LMMs and binomial GLMMs were fitted by maximum likelihood using the functions ‘lmer’ and ‘glmer’ from the ‘LmerTest’ package in *R* (Kuznetsova et al., 2017). Model parameter estimates and their 95% confidence intervals were produced using the ‘stargazer’ package in *R* (Hlavac, 2018). Power analysis was performed using the ‘simr’ package in *R* (Green and Macleod, 2016), with 100 iterations per parameter set. All analyses were performed in *R* (R Core Team, 2023).

### Inference of the genetic basis of sex and sterility determination —

To investigate the genetics of sex determination and male sterility in *M. annua*, we performed crosses between different sexual phenotypes in the greenhouse from 2015 to 2018. Crossed individuals were seeds sampled directly from females in the wild populations. We also continued crosses over two generations for a subset of parents (and grandparents) sampled in 2015. To perform all crosses, we grew the relevant parents, and placed them together in 0.5 L pots. Following our previous breeding protocol, which has shown less than one percent of pollen contamination (Pannell, 1997a), we put each group of crosses sharing the same pollen donor into a pollen-proof growth box into which there was constant filtered airflow to minimize contamination by any outsider pollen. We allowed the parents to mate over the period of a month. For each cross, one pollen donor (either male or hermaphrodite) was mated with one, two or three females. After mating, plants were left to develop fruits and seeds, which were then collected and stored for three months before being germinated and sexed. We restricted our analysis to all families from crosses that generated more than 20 offspring. In total, there were 64 crosses, 40 between a female and a male, 18 between a female and a hermaphrodite, and six self-fertilized hermaphrodites.

We used crosses between females and hermaphrodites (F x H), and hermaphrodites and hermaphrodites (H x H; selfed hermaphrodites) to infer the genetics of male sterility. For these crosses, we restricted our analyses to those generating at least 20 progeny and used Chi-square goodness-of-fit tests (in the *XNomial* package; Engels, 2015) to evaluate evidence for the rejection of a number of plausible simple models for the genetic basis of male sterility. Significance levels were adjusted using the Bonferroni method to account for multiple tests.

We also conducted crosses between females and males (F x M) to assess potential models for the combined effects of male determination and the genetics of male sterility. Here, we included in our evaluation the possibility that male offspring themselves might express male sterility, i.e., these crosses involved four potential progeny classes: males, females, hermaphrodites and sterile males. Due to the greater complexity of the possible genetics underlying progeny ratios from crosses involving males, and because progeny numbers here were lower for each sex category, we analyzed progeny sex ratios using a log-likelihood ratio test using the *XNomial* package (Engels, 2015) in *R*, with P-values calculated for each cross using a resampling procedure (Engels, 2009; Resin, 2023).

Our preliminary analyses pointed to a possible genetic linkage between the male determiner on the Y chromosome of *M. annua* and male fertility restorer loci on the same chromosome; under this model, males carrying a sterility cytotype and one or more fertility restorers would express a normal male sexual phenotype, whereas those lacking a restorer would be male-sterile. To evaluate total evidence for this more complex model, we calculated the log-likelihood of models with different possible recombination rates between the putative male-determination and restorer loci on the Y chromosome. We adopted an approach similar to Fisher’s weighted combined probability method (Fisher, 1936; Zelen and Joel, 1959). Specifically, we multiplied the likelihood ratios of all crosses, weighted by their sample size, according to

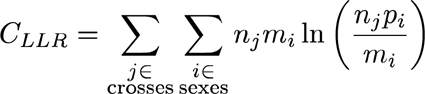

where *C_LLR_* is the log-likelihood ratio of fitting the same genetic model to the results of all F x M crosses, *n_j_* is the number of total offspring from cross *j* (within 39 F x M crosses, one outlier excluded; see Results), *m_i_* is the number of offspring exhibiting sex *i* (male, female, hermaphrodite, male-sterile), and *p_i_* is the probability of that gender according to the hypothesized model. *C_LLR_* is maximized with *m_i_* = *np_i_* for all *i*.

We varied the recombination rate between the male determiner and each of the two putative restorer loci from 0.0 and 0.5 at 0.01 increment and then computed the corresponding value of *p_i_*, and thus the value of *C_LLR_*. We determined the best-fitting recombination rate as that value yielding the greatest value of *C_LLR_*.

Note that for each combination of recombination rates, there are different scenarios of male genotypes depending on the heterozygosity and which allele resides on which sex chromosome (see main text), which result in different expected sex ratios. We calculated the log- likelihood ratio of all these expected sex ratios to the empirical sex ratio of each cross and selected the value of the best-fit scenario as the score of fitting the model to this cross, which then be used to calculate the overall fitting value *C_LLR_* using the formula above.

### Deterministic simulations with inferred parameters —

After estimating the fitness of males and females relative to hermaphrodites and inferring the mode of genetic determination of sex, we performed deterministic simulations (assuming an infinite diploid population) to predict the equilibrium frequencies of the three genders in large populations. We considered a wide range of relative male fitness, from six to 15, based on estimates of Pannell (1997c) and those from this study, a range of relative female fitness from 1.0 to 1.11, based on our estimates here. We considered selfing rates between 0.0 and 1.0 and inbreeding depression between 0.0 and 0.5. Previous work has estimated selfing rates for androdioecious *M. annua* of between 0.1 and 0.3 (Eppley and Pannell, 2007a; Pujol et al., 2009; Korbecka et al., 2011) and inbreeding depression of between 0.0 and 0.5 (Pujol et al., 2009; Pannell et al., 2014).

In our simulations, we assumed that the cost of fertility restoration in terms of both pollen and seed production was *rho* and assumed that the alleles for restoration was fully dominant. Restorer alleles at both restorer loci were assumed to bear the same cost, which we combined multiplicatively between loci. At the start of the simulation, 64 genotypes (nucleus: one XY locus and two nuclear restorer loci; cytoplast: two cytotypes, one male-sterile and one male-fertile) were all present in the population, at arbitrary frequencies. For each generation, the pollen haplotype frequencies were calculated by summing the contributions of each genotype, using its frequency, its pollen production and Mendelian segregation ratios and assumed recombination rate among loci. Ovule haplotypes were calculated similarly, taking inbreeding depression into account. Double crossover between any two considered loci was ignored. Zygote frequencies for the next generation were calculated based on the ovule and pollen frequencies, assuming a combination of selfing and random mating. Evolution of the population was simulated until it reached equilibrium (in practice, when we observed no further change in allele frequencies greater than 10^-6^ over 20 generations). Source code for the simulations, statistical analysis, and raw data will be available upon publication.

## RESULTS

### Spatial and temporal variation in sex ratios ***—***

We surveyed 109 populations of *M. annua* for sexual morph frequencies (Figures 1, 2A; Table S1). We found 36 populations comprised only hermaphrodites, 25 were androdioecious (with males and hermaphrodites), 12 were gynodioecious (with fertile hermaphrodites and hermaphrodites with male sterility, henceforth ‘females’), and 36 were trioecious (with males, female and hermaphrodites). Overall, females, males and hermaphrodites represented 5.3%, 8.4% and 86.3% of all plants observed, but frequencies varied among populations and regions. Females and males were more frequent in the south, along the Mediterranean coast, especially around Gandia (with 18% females and 20% males), Malaga (7.6% and 9.1%) and Valencia (6.3 % and 13%); indeed, most sexually polymorphic populations (gynodioecious, androdioecious, and trioecious populations) were found in these latter regions.

**Figure 1.**
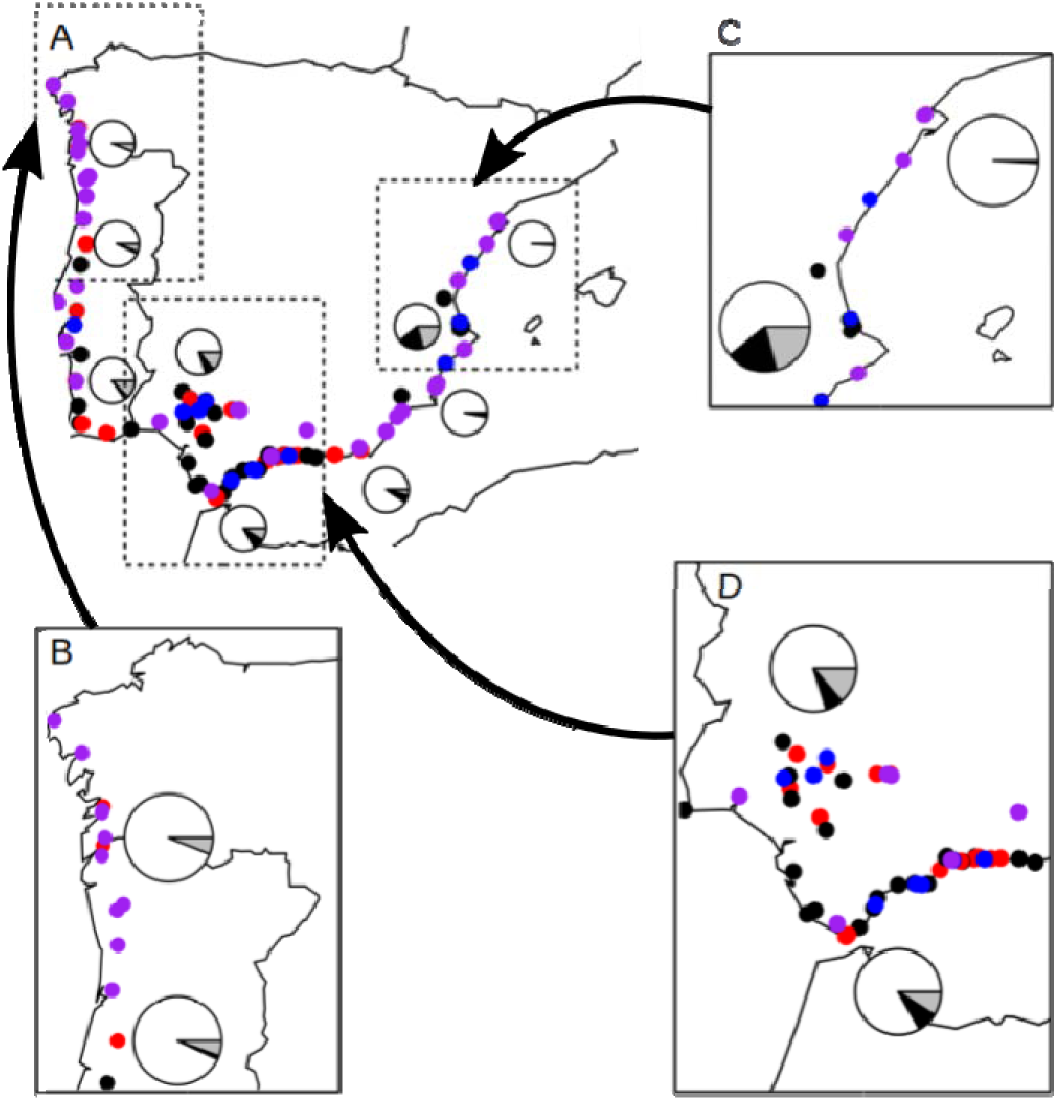
Distribution of the sampled populations of *M. annua* around the Iberian Peninsula as a whole (a), as well as in the north-western (b), eastern (c), and southern (d) Iberian Peninsula. We chose an arbitrary low-frequency cut-off for defining the sexual system of populations visited: populations with > 2.5% females and > 2.5% males were considered trioecious; those with a female frequency > 2.5% but a male frequency < 2.5% were considered gynodioecious; those with a female frequency < 2.5% but a male frequency > 2.5% were considered androdioecious; and those in which the frequencies of both females and males < 2.5% were considered hermaphroditic. Hermaphroditic populations are displayed in purple, androdioecious populations in red, gynodioecious populations in blue and trioecious populations in black. The pie charts represent the frequency of each sex in the area they are close to (black: females; grey: males; white: hermaphrodites).

**Figure 2.**
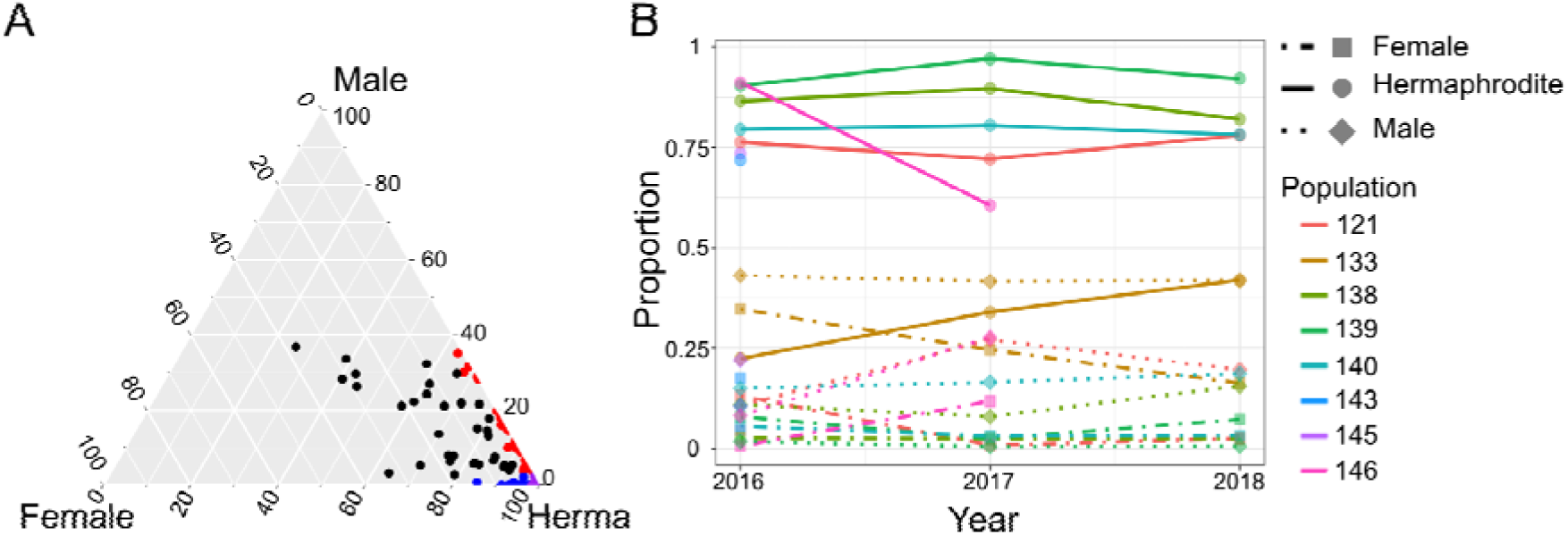
(A) De Finetti diagram of sexual morph frequencies in *M. annua* across the Iberian Peninsula. Black, red, blue, and purple circles represent trioecious, androdioecious, gynodioecious and hermaphroditic populations. See the caption of Figure 1 for the cut-off sex-morph frequency defining each sexual system. (B) The proportions of hermaphrodites (solid lines and round symbols), females (dot-dashed lines and square symbols) and males (dotted lines and diamond symbols) of eight wild populations from 2016 to 2018.

Females were more frequent (mean of 8.2%) in populations with > 2.5% males than in populations with < 2.5% males (1.6%; Wilcoxon test: W = 2182.5, P < 0.0001). Accordingly, females were more frequent in trioecious populations (mean of 13.7%) than in gynodioecious populations (mean of 5%; Wilcoxon test: W = 437, P = 0.003). However, there was no overall correlation between the frequency of females and that of males (binomial GLM: Z = 1.63, df= 106, P = 0.103) nor between the frequency of females and population density (binomial GLM: Z = −0.5, df = 106, P = 0.620).

We recorded the sex ratio in eight populations over three consecutive years (in 2016, 2017 and 2018). Two of these eight initial populations had become extinct in 2017 and a further one had become extinct in 2018, all as a result of human activities. Out of the five persisting populations that were surveyed in both 2016 and 2018, the female frequency decreased in two and remained relatively constant in three, and the male frequency increased in one and remained constant in four (Figure 2B, Table S2). There were no overall directional changes in the sex ratio.

### Allocation to male function by males and hermaphrodites —

In the common garden experiment, we found that males produced more male flowers per biomass than hermaphrodites (LMM: t = 16.7, df = 137, P < 0.001, Figure 3A). The male reproductive effort was 11.7 times higher in males than in hermaphrodites (95% CI: 11.2 - 12.9, Table 1), similar to that estimated previously for a different sample of populations of hexaploid *M. annua* in southern Spain (Pannell, 1997a, c).

**Figure 3.**
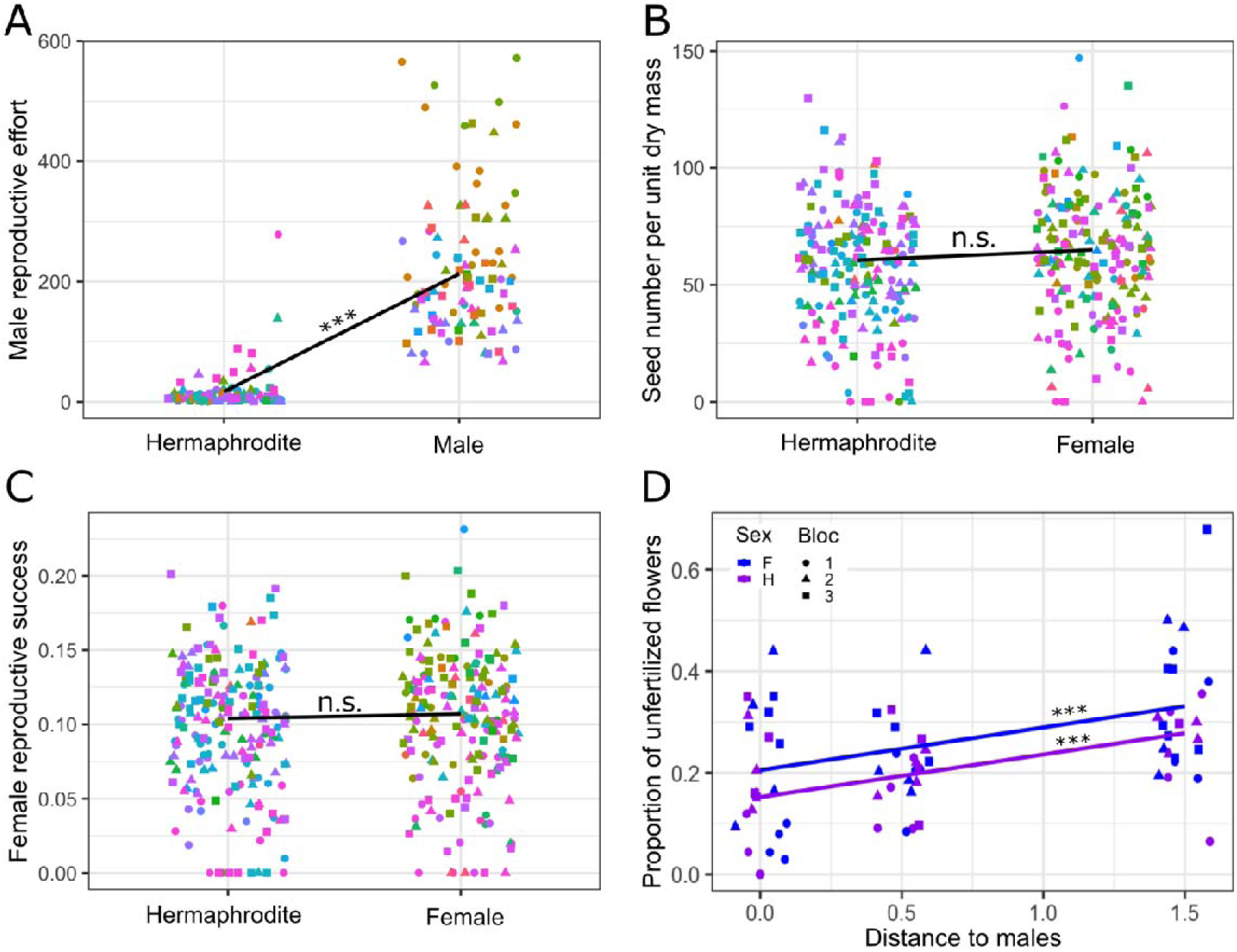
The male reproductive effort of males and hermaphrodites (milligrams of male flower per gram of plant biomass, panel A), the relative female reproductive effort (grams of seed per gram of plant biomass, panel B) and seed number per gram of plant biomass (C) of females and hermaphrodites of *M. annua* measured in the common garden. Colors depict maternal families and shapes depict experimental replicates. Black lines and their slope significance depict the predicted regression lines from LMM models (summary in Table 1). (D) Pollen limitation and distance as the function of the sex. The blue and purple lines represent, respectively, the linear regression predicted by LMM model with sex and distance as fixed factors (Table 2).

**Table 1.**
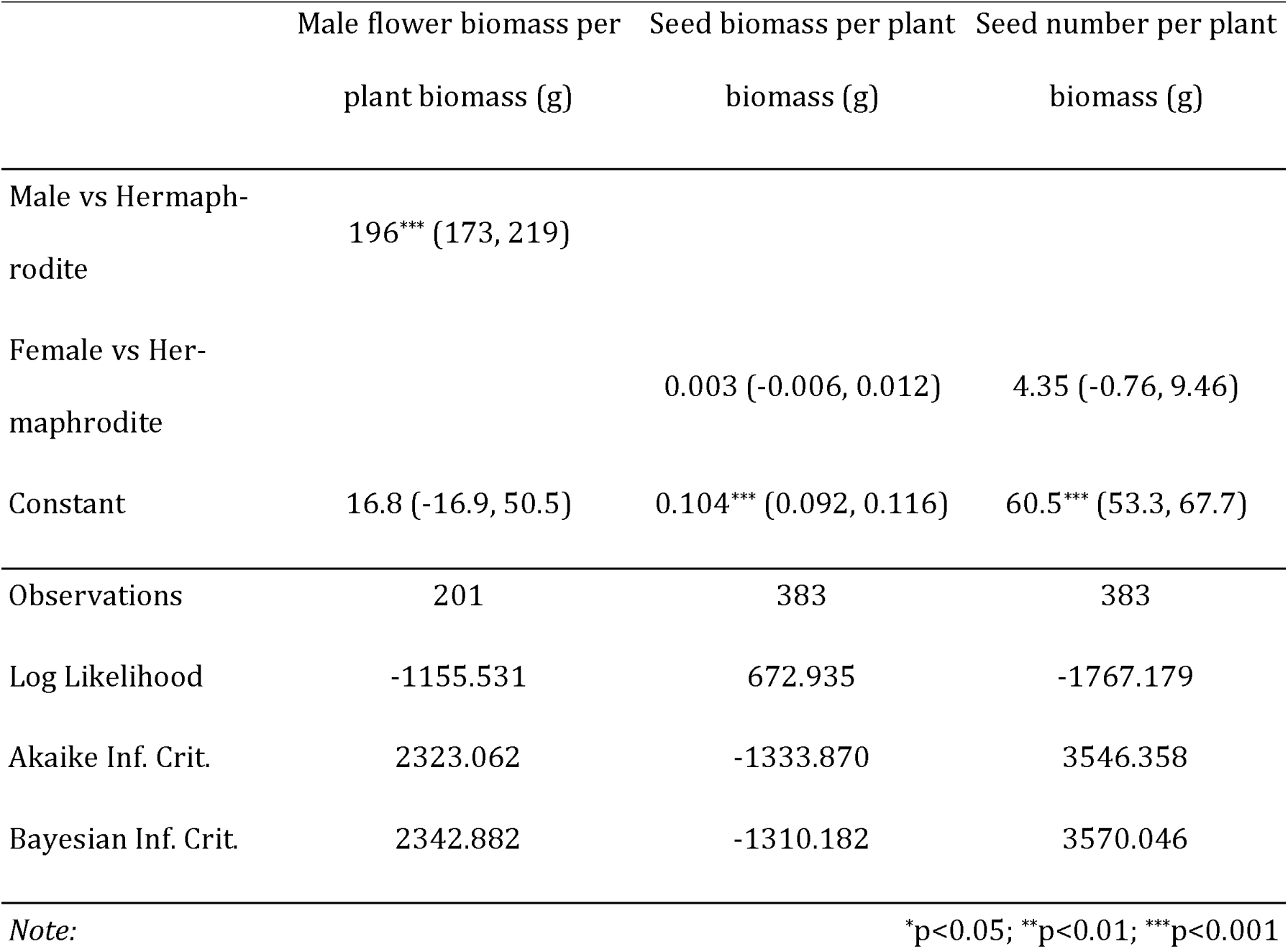
Summary of linear mixed models for relative fitness competition experiment. Confident level = 95%.

**Table 2.** Linear mixed models of the effect of distance from the center and sex on pollen limitation and seed production of female flowers of *M. annua*. Confident level = 95%.

|  | Proportion of unfertilized female flow-<br>ers | Seed biomass per plant bio-<br>mass |
| --- | --- | --- |
| Female vs Hermaphro-<br>dite | 0.053* (0.006, 0.100) | 0.196 (-0.504, 0.895) |
| Distance to male flowers | 0.084*** (0.047, 0.121) | -0.384 (-0.937, 0.169) |
| Constant | 0.152*** (0.072, 0.232) | 3.63*** (2.51, 4.75) |
| Observations | 71 | 71 |
| Log Likelihood | 53.034 | -130.616 |
| Akaike Inf. Crit. | -96.068 | 271.232 |
| Bayesian Inf. Crit. | -84.755 | 282.545 |
| Note: | *p<0.05; **p<0.01; ***p<0.001 |  |

### Allocation to female function by females and hermaphrodites —

Female allocation was similar for females and hermaphrodites, whether measured in terms of reproductive effort (LMM: t = 0.598, df = 213, P = 0.55) or the number of seeds per biomass (LMM: t = 2.906, df = 213, P = 0.097; see Table 1, Figure 3B, 3C). A power analysis suggests that our sampling permitted the detection, with a probability of 0.8, of differences between females and hermaphrodites in female allocation > 12%, irrespective of the measure used, i.e., we can be reasonably confident that females are unlikely to produce greater than 1.12 times the seeds produced by hermaphrodites.

### Estimates of pollen limitation for seed production in females versus hermaphrodites —

Our study of pollen limitation showed that seed production was somewhat pollen-limited for both hermaphrodites and females (LMM with Satterthwaite’s method: t = 4.445, df = 66, P < 0.001), though females were more so (t = 2.228, df = 66.1, P = 0.029; Table 2 and Figure 3D). More specifically, we found that the proportion of unfertilized female flowers increased by approximately 8% per meter’s distance from the central male pollen source, for both females and hermaphrodites (95% CI: 5-12%; Table 2), but for a given distance females had 5% more unfertilized flowers than hermaphrodites (95% CI: 0.6-10%; Table 2), i.e., there was no interaction for pollen limitation between distance and sex (LMM with Satterthwaite’s method: t = 0.245, df = 65.4, P = 0.80, and dropped out of the LMM model based on an AIC comparison). Nevertheless, variation in absolute seed production among individuals overwhelmed the small effects of pollen limitation, so that there were no differences in total seed production between females and hermaphrodites in the experiment (t = 0.548, df = 66.2, P = 0.58), no effect of distance (t = −1.36, df = 66, P = 0.17; Table 2), and no significant interaction (t = −0.613, df = 65.5, P = 0.54, and dropped out of the LMM model based on an AIC comparison).

### Results of crosses —

#### Data used for the analysis of crosses —

We restrict our analysis of data from crosses to all families that generated at least 20 progenies. Of the 64 crosses analyzed, 40 were between a female and a male (F x M), 18 were between a female and a hermaphrodite (F x H) and six were self-fertilized hermaphrodites (H x H). Data are shown in Table S3 and Figure 4.

**Figure 4.**
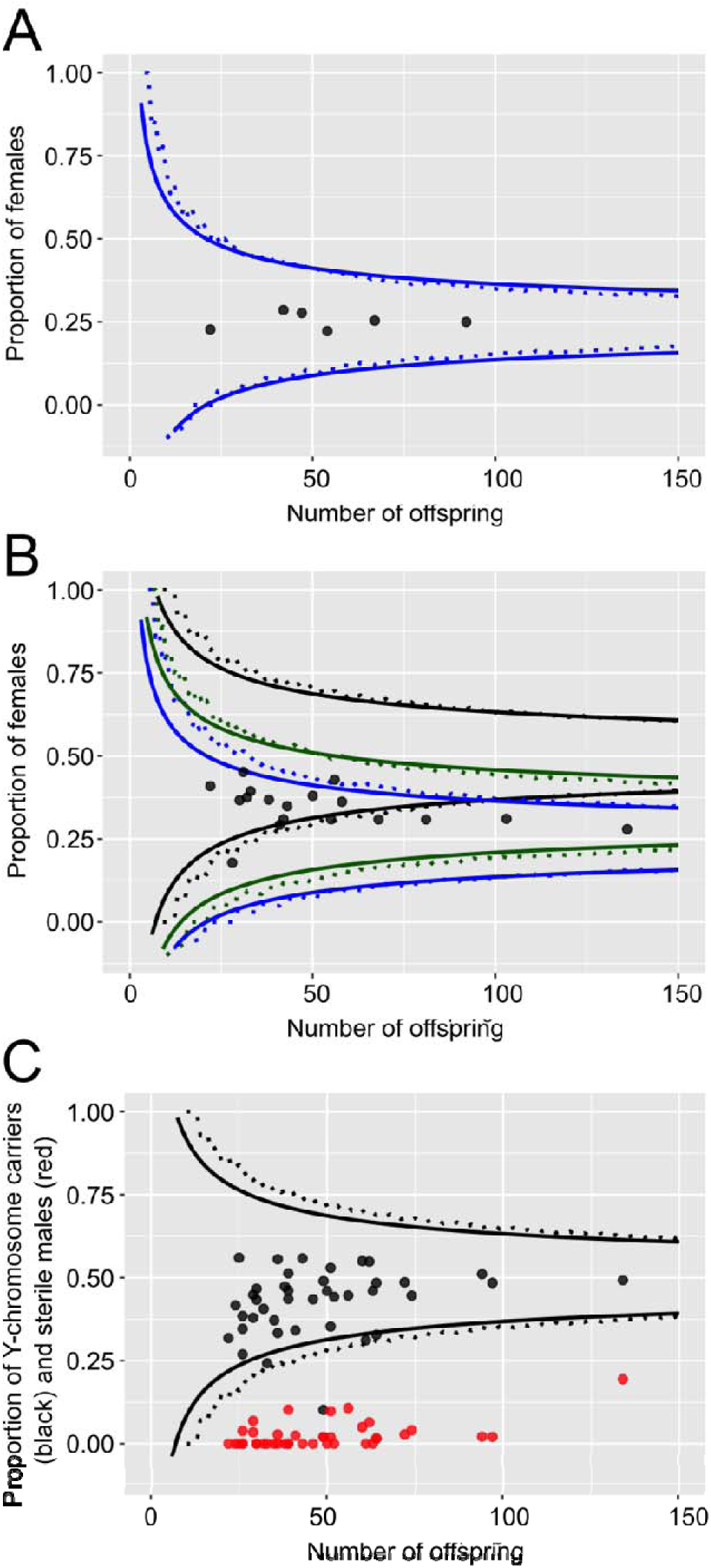
The sex ratios of (A) six hermaphrodite x hermaphrodite crosses (B) 18 female x hermaphrodite crosses, (C) 40 female x male crosses in *M. annua*, plotted against seed family size. Chi-squared acceptance range for each ratio is between the solid lines and the exact binomial probability acceptance range for each ratio is between the dotted lines. (A) The boundaries of the 0.05 (P = 0.05/6, Bonferroni corrected) acceptance region for departures from 1:3 (blue) female-to-the-rest-of-offspring. (B) The boundaries of the 0.05 (P = 0.05/18, Bonferroni corrected) acceptance region for departures from 1:1 (black), 1:3 (blue) and 0.315:0.685 (green) female-to-the-rest-of-offspring. (C) The boundaries of the corrected 0.05 (P = 0.05/40, Bonferroni corrected) acceptance region for departures from 1:1 Y- chromosome-carrier-to-the-rest-of-offspring. Black dots: proportion of Y chromosome carriers (males and sterile males), red dots: proportion of sterile males. The solid lines are the analytical solution, and the dotted lines are the numerical solution. The numerical boundaries show the first cases that were rejected.

#### Model selection and comparisons —

Females were produced in all 64 crosses retained for analysis, whether they involved female parents or not. This indicates that the female determiner (i.e., the male-sterility genes) cannot be attributed to the action of a simple dominant allele segregating at an autosomal locus. The two simplest models that might account for these results thus involve either (1) an autosomal locus with recessive male sterility or (2) an interaction between a cytoplasmic male sterility (CMS) allele and an autosomal restorer allele that would have to be dominant (because selfed hermaphrodites always produce at least some female progeny). We can rule out the first of these two alternatives. Models robustly indicate that females can only be maintained with hermaphrodites with autosomal (or nuclear) male sterility if they produce at least twice the number of seeds as hermaphrodites (Lewis, 1941; Charnov et al., 1976), otherwise, only if inbreeding depression is severe (Charlesworth and Charlesworth, 1978). Yet our above analysis indicates that females produce no more than 1.12 times the seeds produced by hermaphrodites, and self-fertilized progeny of hexaploid *M. annua* in the Iberian Peninsula experience low to intermediate depression (Pujol et al., 2009). Taken together, our overall crossing results and these relative fitness estimates strongly point to a model of CMS. In what follows, we thus assume CMS and seek the simplest model involving potential fertility restoration by nuclear genes to account for further details of our crossing results. We return to this assumption in the Discussion.

#### Crosses involving females and/or hermaphrodites —

As the hexaploidy *M. annua* has a disomic inheritance mode (Obbard, 2004), its nuclear loci number can be estimated with the Mendelian segregation ratio. To estimate the number of loci that might play a role in the restoration of male fertility in trioecious *M. annua*, we adopted an approach that involved seeking the simplest scenario capable of accounting for our crossing results. Clearly, more complex scenarios cannot be ruled out, but we take this approach as a first approximation of the genetics of sex determination in these populations. If we assume the role of only one restorer locus, self-fertilized hermaphrodites would be expected to produce either no females (in the case of a homozygous restorer locus) or 25% females (in the case of heterozygous). Similarly, crosses between a female and a hermaphrodite would be expected to produce either no females (if the restorer locus is homozygous in the hermaphrodite sire) or 50% females (if the restorer locus is heterozygous in the sire). Selfed hermaphrodites always produced about 25% male-sterile individuals in their progeny (Figure 4A), consistent with all of them being heterozygous at a single restorer locus. However, the results of five F x H crosses are inconsistent with a model involving a single restorer locus: all of these five crosses progeny with a strong deficit of females (Figure 4B), with the frequency of females being significantly < 0.5 in four of the five crosses after Bonferroni correction, and the remaining cross being marginally significant.

With the rejection of a one-locus model for fertility restoration, we next considered a scenario involving two independent restorer loci, again assuming dominance for the restoring allele at both loci. A scenario in which hermaphrodites are heterozygous at both restorer loci would yield a frequency of females of 0.25 in female x hermaphrodite crosses, assuming of course that the female mothers are homozygous at both restorer loci for the non-restoring allele. This scenario fits all five of our crosses (Figure 4B in the area between the black-blue lines). Since the two models are not mutually exclusive but the two-loci model is an extension of the one-locus model, all our data from F x H and H x H crosses can therefore be explained by a model involving two restorer loci with dominant restoration at each.

#### Maleness is determined by an XY sex-determination system —

Apart from two exceptions, male progeny were only ever produced in crosses involving a male sire, and all crosses with male sires produced at least some male progeny. These results are fully consistent with single-locus determination of maleness and thus an XY sex determination, as previously inferred for androdioecious *M. annua* (Pannell, 1997a; Russell and Pannell, 2015). The two exceptions involved the production of male progeny in crosses between a female and a hermaphrodite (Table S3, crosses 14.1 x 6.1 and 27.6 x 6.1). We believe that these are likely the result of contamination by pollen from a male in the greenhouse. We repeated the cross 14.1 x 6.1 in 2017 with same-sex siblings of mothers and fathers from those in 2015, with two replicates each (the cross 27.6 x 6.1 could not be repeated due to the lack of 27.6 siblings and disease in the greenhouse). These two replicates showed no male offspring, consistent with the supposition of contamination in the previous cross.

Consistent with an inference of XY determination of maleness, all but one of the 40 crosses between males and females produced sex ratios that did not deviate significantly from 1:1 (Figure 4C). One of the crosses, producing a total of 49 progeny, yielded only four fertile males and one sterile male (see next paragraph), representing a strong deficit of males. We have no explanation for this outlier.

#### The Y chromosome is partially linked to the two CMS restorers —

Some male progeny from our F x M crosses were male-sterile (we refer to these individuals as ‘sterile males’), but these individuals always bore the typical male inflorescence architecture (possession of a pedunculate inflorescence stalk) and are thus unambiguously male; we suppose they carry the Y chromosome and result from the expression of the CMS mutation in the absence of fertility restoration. Hence, F x M crosses may produce: males (with the Y chromosome and a restorer allele at one or both putative restoration loci); sterile males (with the Y chromosome and no restorer allele at either of the two putative restoration loci; females (with no Y chromosome and no restorer alleles at either locus); and hermaphrodites (with no Y chromosome and at one restorer alleles at one or both putative restoration loci). The three loci involved in this phenotypic variation may be segregating independently or may be (partially) linked. We evaluate the possibility of such linkage with our crossing results.

If the Y locus and the male sterility restoration loci are unlinked, two different sex ratios can be expected for F x M crosses, depending on whether the male is heterozygous at one or both restoration loci. A cross between a female and a male carrying one restorer is expected to yield 1:1:2 female-to-hermaphrodite-to-Y-chromosome-carrier. Of 40 F x M crosses, six crosses were inconsistent with this expectation (Chi-squared goodness of fit test, P < 0.001). Among these six crosses, one cross had an excess of hermaphrodites (3:22:8 female-to-hermaphrodite-to-Y-chromosome-carrier, X-squared = 30.6, df = 2), two had an excess of females (29:14:21, X-squared = 14.6, df = 2; and 53:15:66, X-squared = 21.6, df = 2), two had an excess of females and a deficit of hermaphrodites (26:5:25, X-squared = 16.4, df = 2 and 34:7:33, X-squared = 20.6, df = 2) and, finally, one (the one outlier cross referred to above) had a strong deficit of males. Alternatively, if we assume that the male sire is heterozygous (carrying a dominant restorer allele) at both restoration loci, we should expect a sex ratio of 1:3:4 female-to-hermaphrodite-to-Y-chromosome-carrier among the offspring. This ratio is consistent with the results of only one of the above-mentioned crosses (17.1 x 8.2, 3:22:8, X-squared = 12.2, df = 2, P = 0.002). The results of the other five F x M crosses yielded an excess of females and thus remain unexplained by a model involving two restorer loci unlinked to the Y male determiner. We conclude that, with this 3-locus model, independent segregation of alleles can be ruled out. Note that we also tested a model with up to 4 unliked restorer loci, none of these models with a higher number of unliked loci improve the fitting with these five crosses with excess females (data not shown).

Let us now assume that the Y locus and the two fertility restoration loci (R1 and R2) are partially linked on the genetic map in the order R1-Y-R2. We consider this order because we found no evidence for tight linkage between R1 and R2 (the results of all F x H crosses fit to one or two unliked restorer loci, Figure 4B). Ignoring the results of the outlier cross with a major unexplained deficit of males (see above; this cross remains puzzling), and assuming partial R1-Y-R2 linkage, we sought the recombination rates between R1 and Y and between Y and R2 that yielded the best fit to the data of the remaining 39 F x M crosses. We found that the best-fit recombination rates, respectively, were 0.11 and 0.26 (sum of weighted log-likelihood ratio = −3.54). No F x M crosses were statistically inconsistent with this model (Table S4), with 29/39 crosses fitting best to the scenario that the male sire carries both restorers on the Y chromosome (Figure 5B). Furthermore, given this best-fit genetic map, we estimate a recombination rate of 0.37 between R1 and R2 (with the male-determining locus between them on the Y chromosome). Under this scenario, an F x H cross would produce 31.5% females if the hermaphrodite sire carries a single restorer allele at both loci. Our data from the F x H crosses indeed fit better to this scenario than one assuming no linkage (Figure 4B, green lines). Overall, this genetic model, with the male-determining locus situated between two restorer loci on the same (Y) chromosome under partial linkage is capable of explaining all our crosses in all three categories (H x H, F x H and F x M).

**Figure 5.**
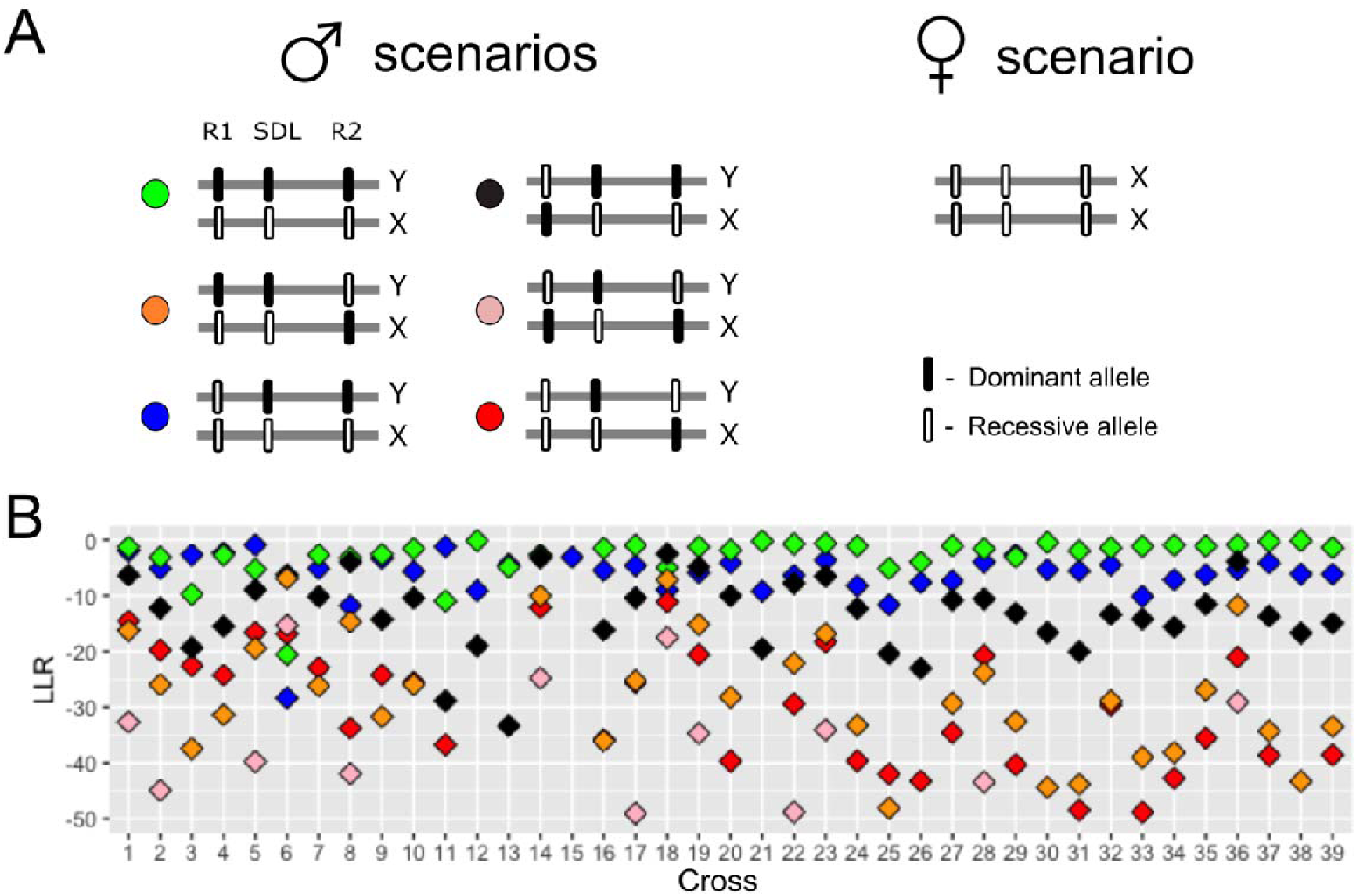
Results of the best-fitting model for the inferred scenario of sex-morph determination for trioecious *M. annua,* in which a single sex-determining locus (SDL) with dominant male determination is flanked on either side by a male-fertility restorer locus segregating for a dominant restorer and a recessive non-restorer (loci R1 and R2, respectively). Our data were most consistent with a genetic map with recombination rates of 0.11 and 0.26 for R1-SDL and SDL-R2, respectively. (A) Depiction of the six best-fit scenarios of sex-chromosome genotypes that could emerge from the assumed base model. Each scenario is color-coded and varies in the combination of the dominant or recessive alleles at each locus of the three sex-morph loci on the X and Y chromosomes. (B) The log-likelihood ratios (LLR) obtained using the *XNomial* package in *R* to evaluate the fit of each of the different scenarios depicted in (A); symbol colors in (B) refer to the respective color labels in (A). Sample sizes for each cross and the P*-*values for the respective LLR tests are presented Table S4. See main text for further details.

### The sex ratios from deterministic models with inferred fitness values and genetic architecture —

Our deterministic simulations indicate that trioecy is possible with this genetic architecture, at the estimated low female fitness of < 1.12 and high male fitness of 12, without the need for pollen limitation (Figure 6). Trioecy can only be maintained in this deterministic simulation when the restorer carries a cost > 0 (Figure 6A). This result conforms with the results of previous work on the maintenance of gynodioecy, which showed that CMS cannot coexist with the male fertile cytotype unless there is a cost of restoration (Charlesworth and Ganders, 1979; Charlesworth, 1981; Delannay et al., 1981). The cost of restoration has been found in multiple species with CMS and is well-studied theoretically (Charlesworth, 1981; Delannay et al., 1981; De Haan et al., 1997; Wen and Chase, 1999; Bailey, 2002; Bailey et al., 2003; Dufaÿ et al., 2007, 2008; Montgomery et al., 2014). Assuming a cost of restoration, our simulations yielded sex ratios similar to those observed for *M. annua* in the field, i.e., with hermaphrodites, males, females and male-sterile males occurring at frequencies between 60 and 100%, 0 and 40%, 0 and 40%, and less than 1%, respectively. This range of sex ratios requires a cost of restoration between 0.15 and 0.4, a selfing rate between 0.3 and 1.0, and inbreeding depression between 0.1 and 0.5 (Figure 6B). If we assume lower selfing rates (0 to 0.3; (Eppley & Pannell, 2007a; Pujol et al., 2009; Korbecka et al., 2011), our simulations predict higher male frequencies than typically observed in the field (Figure 6C). Populations with lower inbreeding depression lose their females (Figure 6D).

**Figure 6.**
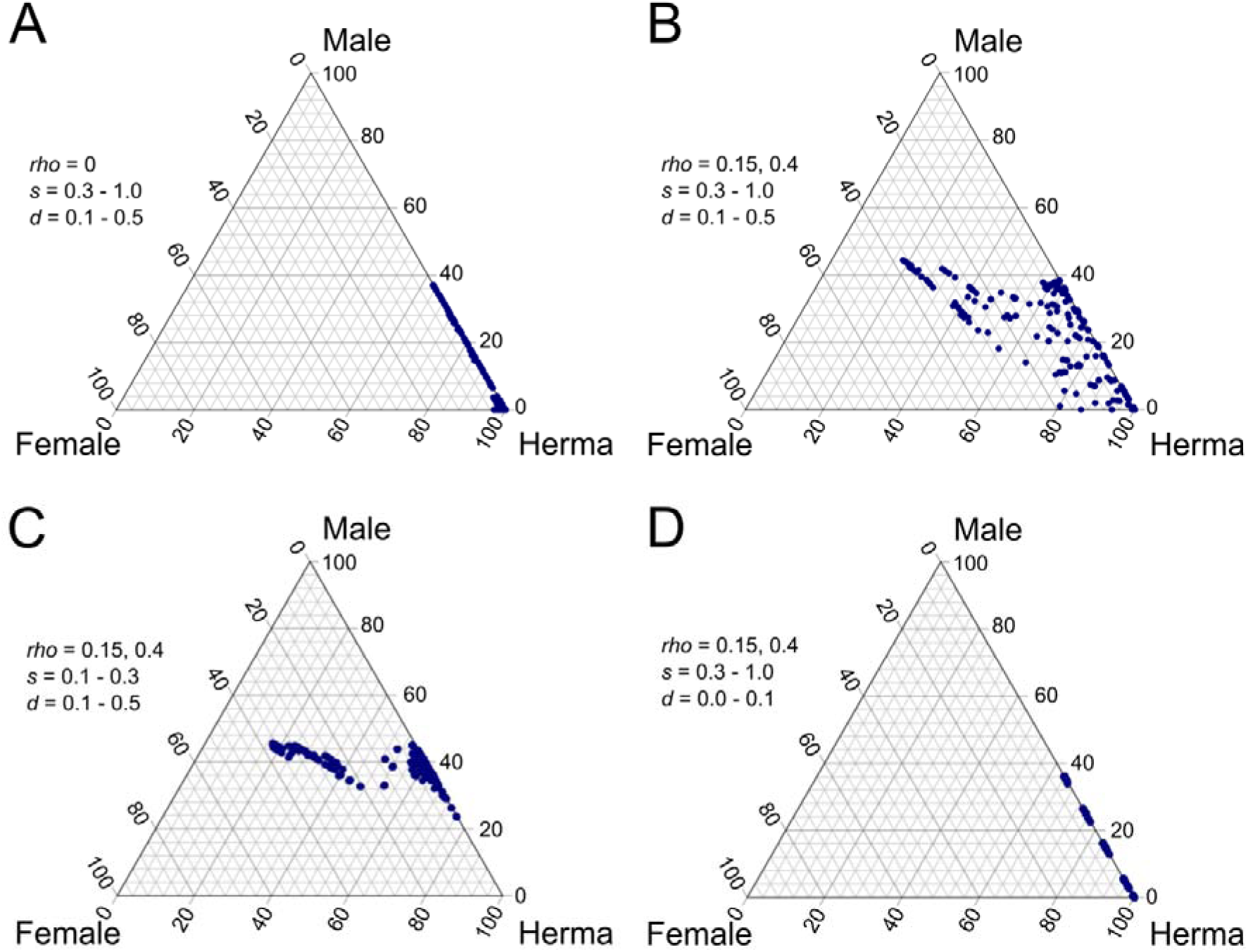
De Finetti diagram of equilibrium sex morph frequencies in simulated populations, with maleness assumed to be determined by alleles at a sex-determining locus (SDL) and femaleness determined by two recessive loci linked to the SDL, with recombination rates of 0.11 and 0.26, as inferred from our model fitting for data from crossed with *M. annua* (see Figure 5A for a depiction of loci locations). Simulations further assumed that males produced between 6.0 and 15 times more pollen than hermaphrodites, and that females produced between 1.0 and 1.11 times more seeds than hermaphrodites. Values assumed for the selfing rate *s*, inbreeding depression *d*, and the cost of restorers *rho* are indicated in the inset legends. ‘Male’, ‘Female’ and ‘Herma’ refers to males, females and hermaphrodites.

## DISCUSSION

### Widespread trioecy in hexaploid M. annua —

Our results indicate that trioecy is widespread in hexaploid *M. annua* in the Iberian Peninsula. Previously, Perry et al. (2012) found evidence for trioecy in several populations in southeastern Spain, but the same three phenotypes co-occur in populations of the species throughout much of its Iberian range, in about 1/3 of the populations we sampled, with a somewhat greater prevalence in the south. The presence and frequency of the three phenotypes vary substantially among populations, many of which were monomorphic (comprising hermaphrodites) or dimorphic (either androdioecious or gynodioecious). Sex ratio variation in hexaploid *M. annua* has previously been described in the context of androdioecy in a number of studies (Pannell, 1997a, b, c; Pannell et al., 2008, 2014), with no reference to the male-sterile phenotypes documented here. In those studies, hermaphrodites will undoubtedly have included both fertile and male-sterile phenotypes, such that some of the populations labeled androdioecious were probably trioecious in the sense we interpret them here. Females in trioecious populations of *M. annua* are phenotypically monoecious, producing sterile male flowers that are otherwise identical to those of fertile hermaphrodites and thus easy to overlook. We might thus characterize trioecy in *M. annua* as the outcome of the invasion and maintenance of male sterility in an androdioecious context, with populations that lack males likely having been colonized by one or a few hermaphrodites in a broader metapopulation (Pannell, 1997b, 2000). The presence or absence of females in hexaploid *M. annua* might thus similarly be interpreted in terms of the outcome of colonization and migration dynamics in a metapopulation. In other words, much of the variation in sex ratios documented here likely has a stochastic origin.

### Relative male and female fertilities of the three sexual phenotypes of trioecious *M. annua**—***

Our results confirm previous work on androdioecious *M. annua* that males produce and disperse much more than twice the amount of pollen than hermaphrodites (Pannell, 1997c) – approximately 12 times as much based on our estimates of male flower production in the present study. Assuming that relative pollen production is closely aligned with relative siring success (Gregorius et al., 1987; Holsinger, 1991), as seems reasonable for a wind-pollinated plant, these estimates of relative male fitness are fully consistent with theoretical predictions for the maintenance of males with hermaphrodites (Charlesworth and Charlesworth, 1978; Charlesworth, 1984, 1999).

In contrast to our estimates of relative male fitness for males, our estimates based on the seed production of females versus hermaphrodites indicate that the female component of reproductive success of females is almost certainly < 1.12 times that for hermaphrodites. This is thus substantially lower than the twofold threshold required for the maintenance of females with biparentally inherited male sterility in fully outcrossing populations (Lewis, 1941; Lloyd, 1975; Charnov et al., 1976). Theory indicates that biparentally inherited male sterility can be maintained in hermaphroditic populations when females have a relative seed production < 2 if hermaphrodites self-fertilize their progeny and inbreeding depression is severe. However, selfing rates in hexaploid *M. annua* tend to be low, especially dense populations and especially in the presence of males (Eppley and Pannell, 2007a; Dorken and Pannell, 2008; Korbecka et al., 2011), and inbreeding depression tends also to be low (though intermediate levels have been estimated for the southern Iberian Peninsula) (Pujol et al., 2009). Taken together, these results strongly suggest that the relative seed production of females is too low to account for their maintenance under a model of biparental inheritance of male sterility. We thus suppose that male sterility in hexaploid *M. annua* has a cytoplasmic basis, as is frequently the case in flowering plants (Kaul, 1988).

### The genetic architecture of trioecy in hexaploid *M. annua* —

Our crossing results confirm that males are determined by a Y chromosome and that females and hermaphrodites lack a Y, as inferred previously (Pannell, 1997a; Russell & Pannell, 2015). Interestingly, although the Y chromosome appears to have had a single origin at the base of the genus *Mercurialis*, the Y chromosome in androdioecious and trioecious populations of hexaploid *M. annua* is derived from a copy that introgressed into these populations from an ancestor of the more distantly related perennial species *M. elliptica*, presumably following the Y chromosome’s loss during the evolution of allopolyploid *M. annua* (Gerchen et al., 2022).

If we assume a cytoplasmic transmission of male sterility, as justified in the previous section, our results reject the simplest genetic models and require us to invoke an interaction between cytoplasmic male sterility (CMS) and fertility restorer alleles transmitted biparentally at more than one locus. This complexity, too, conforms to genetic models invoked for the expression and inheritance of male sterility in a large number of other species of flowering plants (Bailey, 2002; Van Damme et al., 2004; Montgomery et al., 2014; Toriyama, 2021). Indeed, Ross (1969) similarly invoked two restorer loci with redundant expression in gynodioecious *Plantago lanceolata*, i.e., for which a restorer at either one or both loci was necessary and sufficient to restore male fertility to individuals carrying the sterility cytotype. While our results thus appear to conform to a model similar to one proposed for other gynodioecious species, as far as we are aware our study is the first to invoke CMS for the maintenance of trioecy.

Unlike previous genetic models for CMS in populations in which males are absent, a plausible model fitting our data for trioecious populations of *M. annua* requires that the two restorer loci we invoke are located in partial linkage on either side of the sex-determining locus on the sex chromosomes, i.e., in each of two flanking pseudoautosomal regions. This hypothesis of linkage between sex-determining and restorer loci is necessary to explain the fertility of males carrying a CMS mutation, and the observation of occasional sterile males requires that the linkage be strong but incomplete. While this model might seem overly complex and arbitrary, the linkage between a sex-determining locus and fertility restorer loci has previously been shown to be necessary in models for the evolution of dioecy from gynodioecy due to CMS (Ross, 1978; Maurice et al., 1993; Schultz, 1994). The plausibility of our model for trioecious *M. annua* is perhaps further strengthened by the reasonably good fit of its prediction for the distribution of sex ratios under the conditions assumed, based on computer calculations, and those observed among the 109 populations sampled in the field. Nevertheless, we note that the validity of a model involving linkage between sex-determining and restorer loci has never been demonstrated by molecular developmental analysis for any species, and the hypothesis requires verification in *M. annua*. As for previous studies of the maintenance of CMS (Charlesworth, 1981; Delannay et al., 1981), our model also requires that fertility restoration is costly, and this, too, still needs to be demonstrated empirically. Further crosses with independent families and more precise fitness estimates for the relevant genotypes would also be valuable, especially as all our crosses were between individuals produced by open-pollinated females in the field (and none were from open-pollinated hermaphrodites).

### Trioecy in the context of sexual-system variation in *M. annua* more broadly —

The observation of widespread trioecy in hexaploid *M. annua* adds further complexity to the remarkable variation in sexual systems documented in previous studies of the species complex, which includes dioecy, subdioecy, androdioecy and monoecious hermaphroditism (Pannell et al., 2014; Cossard and Pannell, 2019). This variation has hitherto been interpreted in terms of the outcome of the balance between frequency-dependent selection on the sex ratio and sex allocation within populations, largely favoring the maintenance of separate sexes, and stochastic loss of unisexuality and the promotion of bisexuality by selection for reproductive assurance during colonization in metapopulations or through range expansions (Pannell, 1997b, 2000; Pannell et al., 2014). Some of this variation in sex allocation and sexual systems is associated with a mosaic of geographic variation in the abundance of populations and inferred rates of metapopulation turnover (Eppley and Pannell, 2007b), which also appear to have a north-south clinal component in the Iberian Peninsula (Dorken et al., 2017). In this context, it is interesting that we have found trioecy in *M. annua* to be more common in southern Spain than further north. It remains to be seen whether this relates in any way to the documented cline of decreasing inbreeding depression from the south to the north (Pujol et al., 2009), or to the (likely related) history of range expansion towards the north (Obbard et al., 2006; see also Pujol and Pannell, 2008) but both factors may have played a role in influencing the distribution of male sterility in hexaploid *M. annua*. Crosses of genotypes sampled in the north, where CMS is rare or absent, with those in the south, might reveal the extent to which, for example, range expansion has shaped the distribution of cytoplasmic genetic diversity in addition to its effects on neutral genetic diversity (Obbard et al., 2006) and diversity at viability loci (Pujol and Pannell, 2008; Pujol et al., 2009).

The observed occurrence of females in trioecious populations of *M. annua* in the Iberian Peninsula is similar to the occurrence of hermaphrodites with exceedingly low (or zero) male allocation in its populations near Fez in Morocco (Durand, 1963; Dorken and Pannell, 2008). In both of these situations, we observe a tendency towards greater separation of the sexes, perhaps reflecting an evolutionary path to dioecy. However, because females expressing the hypothesized CMS in trioecious populations continue to produce male flowers that are simply rendered sterile, whereas those around Fez are hermaphrodites with an extremely low production of male flowers along a sex-allocation continuum, the two scenarios are developmentally different. They may thus exemplify the convergent evolution towards separate sexes via two different paths: one likely involving genetic variation in sex allocation at nuclear loci (in Morocco, but also more generally in hexaploid *M. annua*); and the other involving the possible spread of selfish cytoplasmic elements, which previous work has shown can precipitate the evolution of dioecy when males invade gynodioecious populations (Maurice et al., 1994; Schultz, 1994). If so, this scenario of convergent evolution would join that invoked for the independent evolution of a male-like inflorescence architecture with improved pollen dispersal in two different cryptic species of hexaploid *M. annua* (Santos del Blanco et al., 2019).

### Wider implications and concluding remarks —

The occurrence of individuals with both male and female functions is relatively common in dioecious flowering plants in general, but the ‘hermaphrodites’ in such cases are almost always the result of the expression of inconstant sex expression in males or females (or in both sexes) (Ehlers and Bataillon, 2007; Cossard and Pannell, 2019). This situation also applies to the androdioecious populations of *M. annua* near Fes in Morocco mentioned above, where females represent hermaphrodites at the extreme end of a continuum from substantial to zero male flower production. The distribution of sex allocation in the populations of *M. annua* with males, females and hermaphrodites described in the present study similarly represent a continuum in which males co-occur with hermaphrodites that vary in their sex allocation. However, they qualify meaningfully as trioecious because females are discretely (and developmentally) different from fertile hermaphrodites by virtue of their production of sterile male flowers. Our study thus confirms the previous documentation of trioecy in *M. annua* by Perry et al. (2012) and adds substantially to our knowledge of its geographic extent, its functional implications for fitness, and its possible genetic basis. It thus adds an exemplar to the very few reasonably well-characterized cases of the sexual system. While trioecy clearly can evolve and be maintained, it remains exceedingly rare, and its occurrence in *M. annua* is linked to its likely evolution in androdioecious populations, a sexual system that is also very rare (Darwin, 1871; Charlesworth, 1984; Pannell, 2002). In this sense, the remarkable variation in sexual systems in *M. annua* remains exceptional, and the discovery of trioecy does not strongly shift the conclusions reached previously about its rarity and its unlikely evolution.

Our study nevertheless raises questions about the role CMS might have played in evolutionary transitions between combined and separate sexes in plants. Previous modelling has drawn attention to the evolution of dioecy from functional hermaphroditism via the initial spread of CMS (Ross, 1978; Maurice et al., 1993; Schultz, 1994). To our knowledge, although gynodioecy with CMS is very common in plants, we still have little evidence for it having played a role in the evolution of fully separate sexes along the lines predicted by the models. Given the inference of transitions between monoecious hermaphroditism and dioecy in the evolution of the *M. annua* species complex, our results would seem to identify it as a potential example of the role of CMS in the evolution of dioecy. However, male sterility remains absent or at low frequency in populations across the range of hexaploid *M. annua*, and any tendency towards dioecy in these populations rather involves quantitative shifts in sex allocation rather than the expression of male sterility. Despite decades of research on male sterility and its role in the evolution of dioecy from hermaphroditism, our understanding remains surprisingly poor, particularly with respect to transitions from gynodioecy to dioecy (Spigler and Ashman, 2012) but also with respect to potential transitions involving CMS in particular.

## ACKNOWLEDGEMENTS

We thank numerous undergraduate assistants and Aline Revel for their help in setting up experiments and measuring plant phenotypes and the Swiss National Science Foundation (Grant 31003A_163384) and the University of Lausanne for funding.

## AUTHOR CONTRIBUTIONS

Conceived and designed the project: JRP; conducted fieldwork in Spain and experiments in Lausanne: TM; analyzed and interpreted the data: MTN; conducted computer simulations: MTN; prepared figures and tables and wrote the first draft of the manuscript: MTN and TM; revised and wrote the final version of the manuscript: MTN and JRP. All authors approved the final version of the manuscript.

## Supplementary Information

**Table S1.**
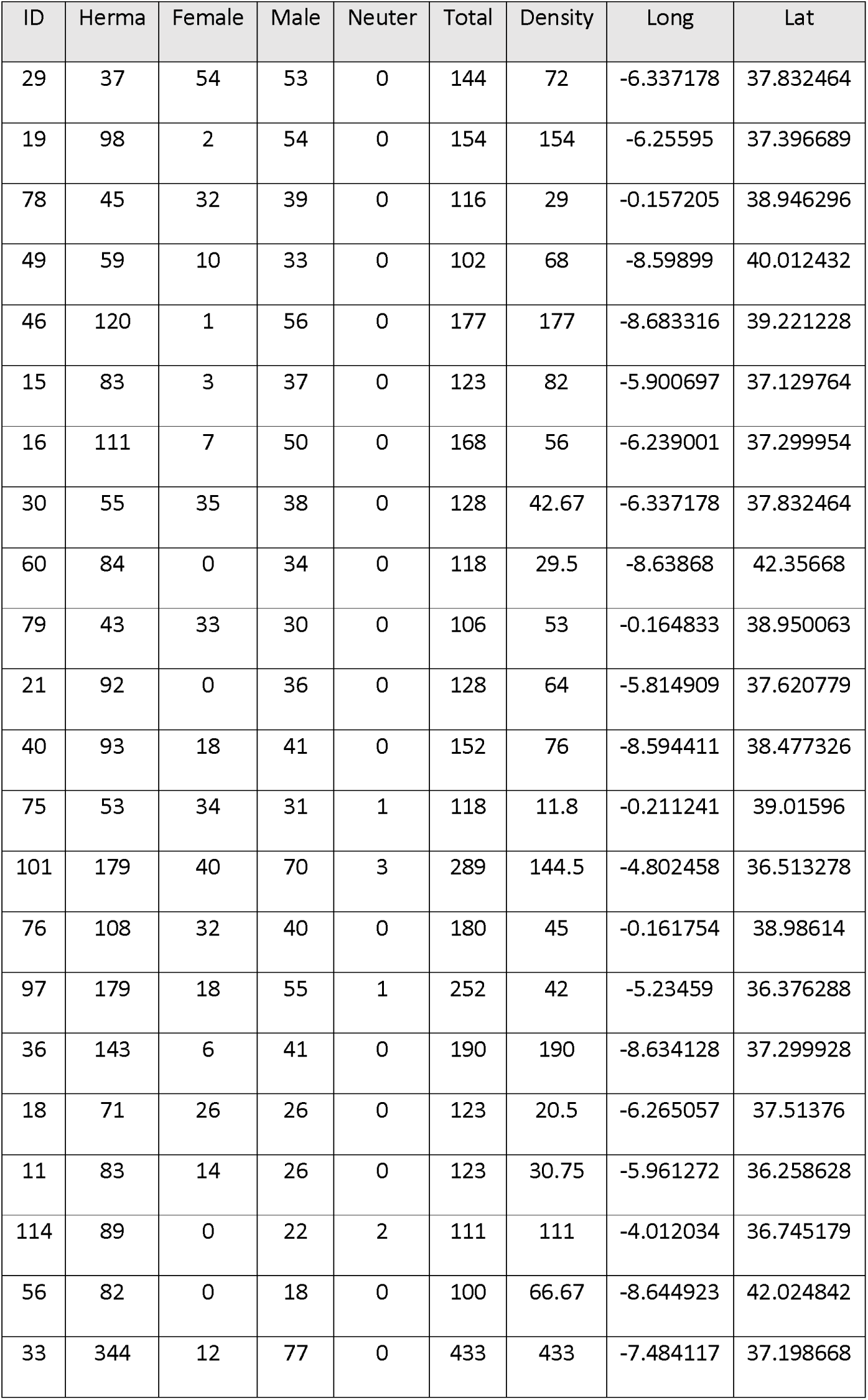
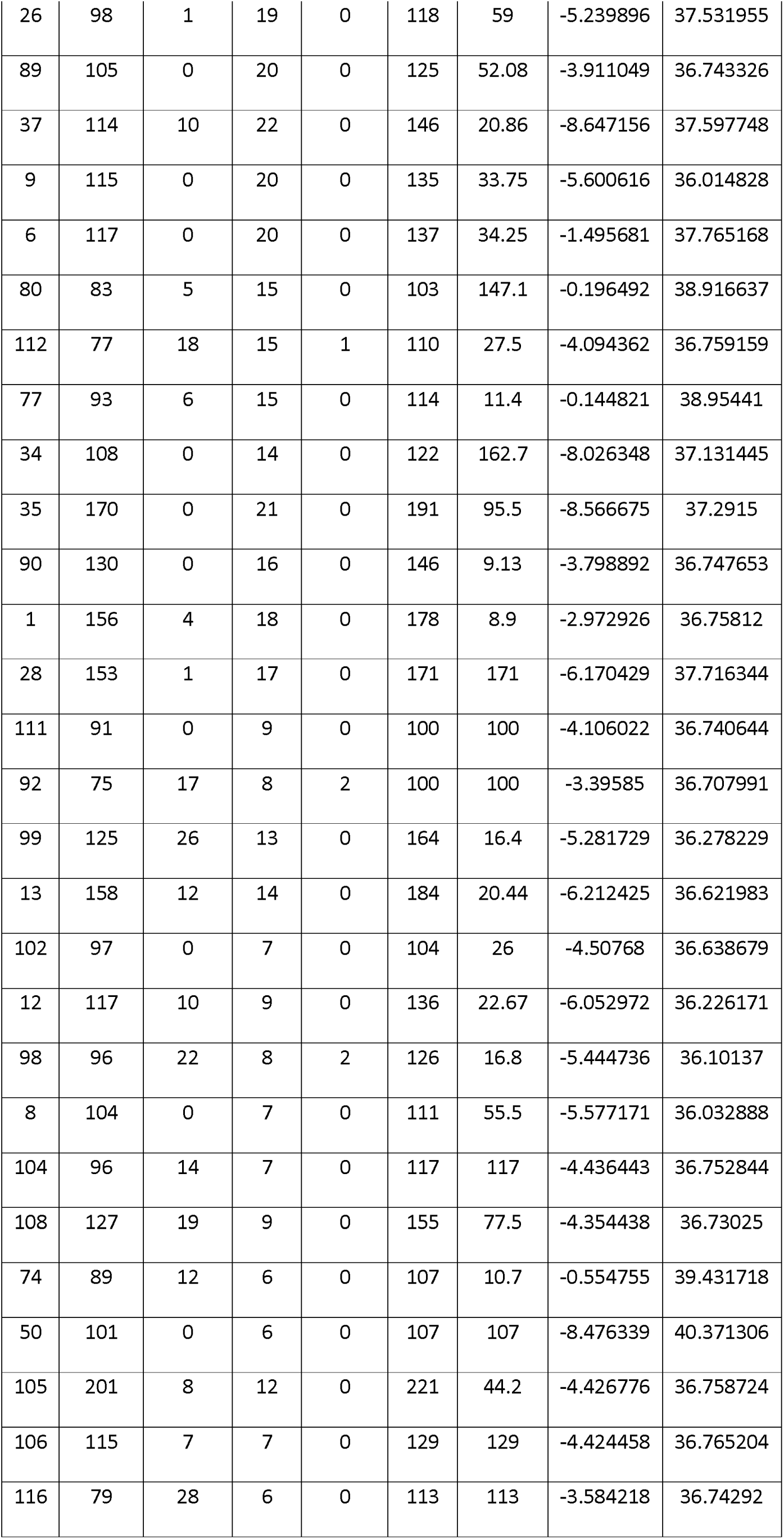
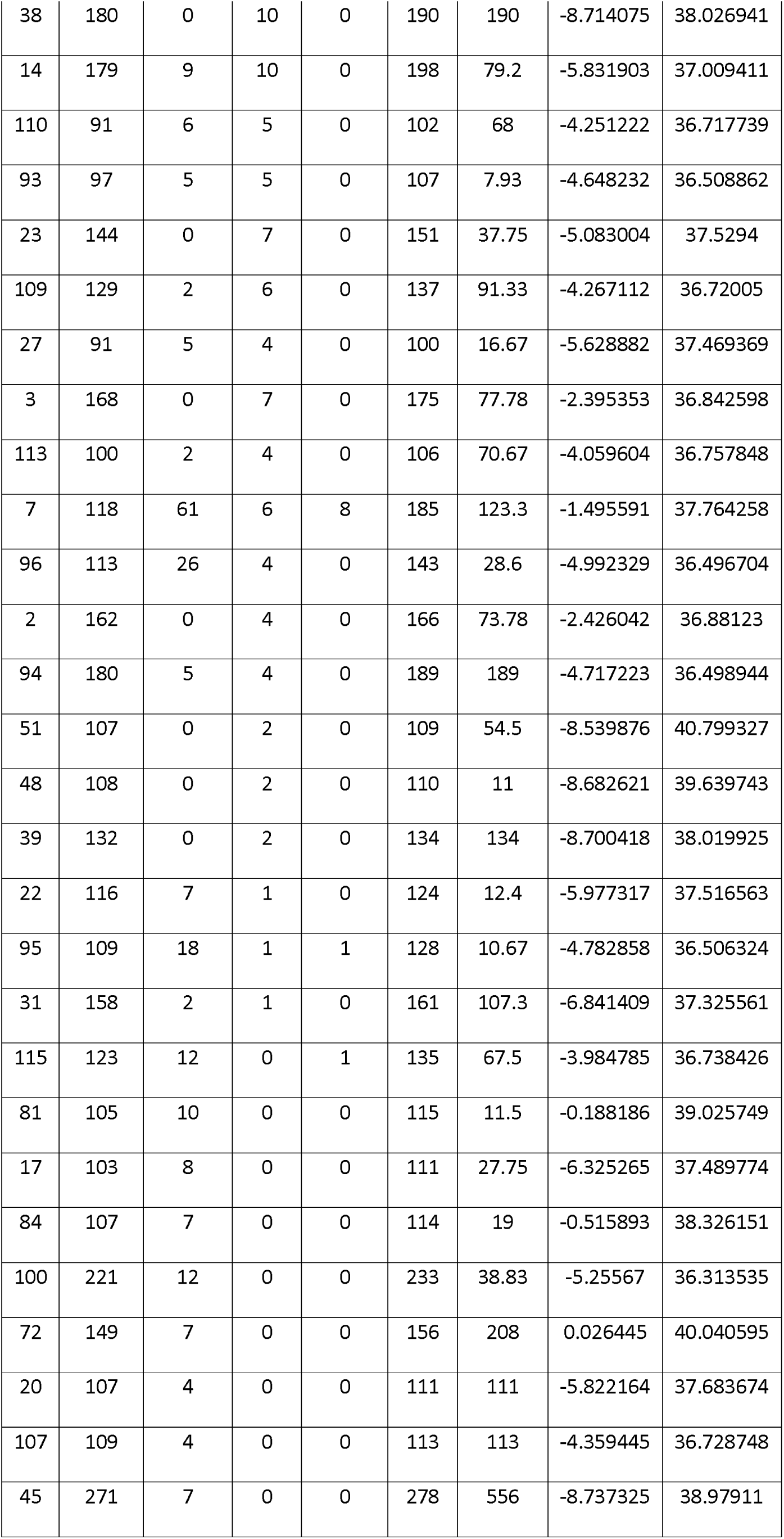
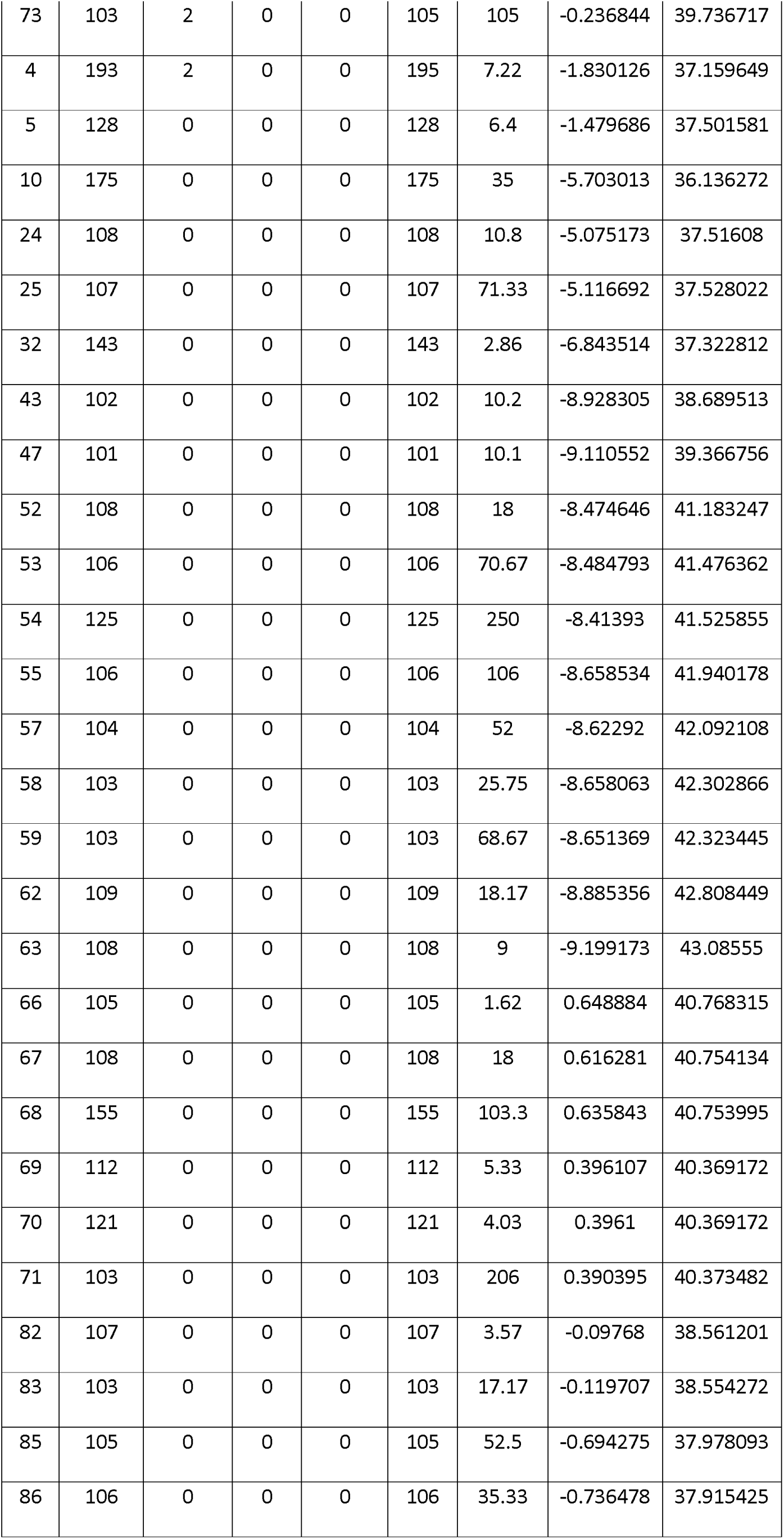
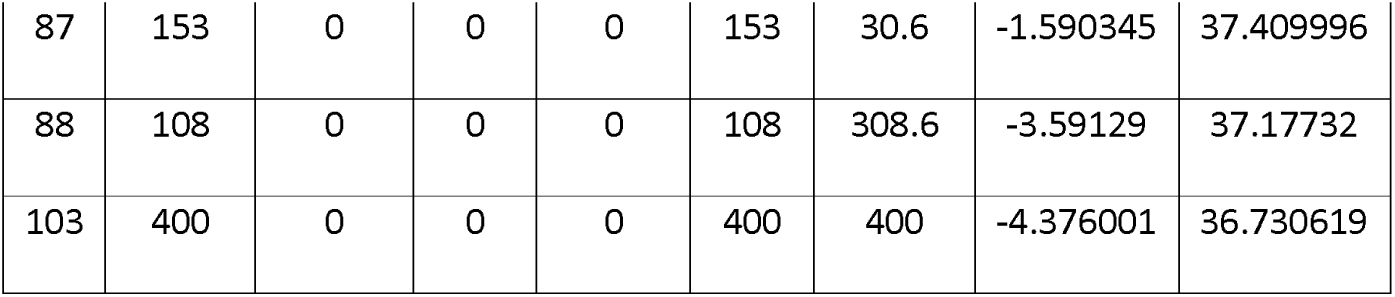
Coordinates, numbers of sexual phenotypes, and density of 109 populations transected in Spain.

**Table S2.** The number of individuals of each sexual phenotype and the density of 8 populations were monitored from 2016 to 2018.

| Population | Herma | Female | Male | Total | Density | Prop_H | Prop_F | Prop_M | Year | GPS |
| --- | --- | --- | --- | --- | --- | --- | --- | --- | --- | --- |
| 133 | 225 | 349 | 434 | 1008 | 42.442 | 0.223 | 0.346 | 0.431 | 2016 | 36.72791,<br>-3.53112 |
| 145 | 361 | 22 | 108 | 491 | 28.045 | 0.735 | 0.045 | 0.22 | 2016 | 36.73339,<br>-3.56282 |
| 138 | 475 | 14 | 60 | 549 | 33.068 | 0.865 | 0.026 | 0.109 | 2016 | 36.75472,<br>-3.88905 |
| 139 | 206 | 18 | 4 | 228 | 12.258 | 0.904 | 0.079 | 0.018 | 2016 | 36.74499,<br>-4.04198 |
| 121 | 180 | 31 | 25 | 236 | 34.483 | 0.763 | 0.131 | 0.106 | 2016 | 36.75723,<br>-4.04596 |
| 140 | 1361 | 95 | 256 | 1712 | 18.708 | 0.795 | 0.055 | 0.15 | 2016 | 36.73672,<br>-3.95346 |
| 143 | 542 | 132 | 80 | 754 | 18.708 | 0.719 | 0.175 | 0.106 | 2016 | 36.73307,<br>-3.51005 |
| 146 | 145 | 1 | 13 | 159 | 47.111 | 0.912 | 0.006 | 0.082 | 2016 | 36.74682,<br>-3.56303 |
| 133 | 704 | 509 | 864 | 2077 | 24.453 | 0.339 | 0.245 | 0.416 | 2017 |  |
| 145 | NA | NA | NA | NA | NA | NA | NA | NA | 2017 |  |
| 138 | 824 | 22 | 73 | 919 | 53.239 | 0.897 | 0.024 | 0.079 | 2017 |  |
| 139 | 204 | 5 | 1 | 210 | 5.833 | 0.971 | 0.024 | 0.005 | 2017 |  |
| 121 | 88 | 1 | 33 | 122 | 16.542 | 0.721 | 0.008 | 0.27 | 2017 |  |
| 140 | 1111 | 42 | 227 | 1380 | 12.477 | 0.805 | 0.03 | 0.164 | 2017 |  |
| 143 | NA | NA | NA | NA | NA | NA | NA | NA | 2017 |  |
| 146 | 408 | 79 | 186 | 673 | 37.444 | 0.606 | 0.117 | 0.276 | 2017 |  |
| 133 | 816 | 316 | 813 | 1945 | 18.524 | 0.42 | 0.162 | 0.418 | 2018 |  |
| 145 | NA | NA | NA | NA | NA | NA | NA | NA | 2018 |  |
| 138 | 540 | 16 | 102 | 658 | 37.443 | 0.821 | 0.024 | 0.155 | 2018 |  |
| 139 | 1235 | 97 | 8 | 1340 | 31.929 | 0.922 | 0.072 | 0.006 | 2018 |  |
| 121 | 318 | 10 | 80 | 408 | 32.64 | 0.779 | 0.025 | 0.196 | 2018 |  |
| 140 | 902 | 37 | 215 | 1154 | 10.414 | 0.782 | 0.032 | 0.186 | 2018 |  |
| 143 | NA | NA | NA | NA | NA | NA | NA | NA | 2018 |  |
| 146 | NA | NA | NA | NA | NA | NA | NA | NA | 2018 |  |

**Table S3.** The number of males, sterile males, females and hermaphrodites in the offspring of 64 crosses.

| Mother | Father | Cross | F | H | M | N | Total |
| --- | --- | --- | --- | --- | --- | --- | --- |
| 10.1 | 24.1 | FxM | 10 | 6 | 9 | 1 | 26 |
| 10.1 | 8.3 | FxM | 13 | 6 | 7 | 0 | 26 |
| 13.2 | 8.3 | FxM | 22 | 11 | 13 | 5 | 51 |
| 14.1 | 6.1 | FxH | 14 | 10 | 6 | 1 | 31 |
| 16.1 | 24.1 | FxM | 15 | 3 | 10 | 1 | 29 |
| 16.1 | 8.2 | FxM | 13 | 8 | 14 | 4 | 39 |
| 17.1 | 8.2 | FxM | 3 | 22 | 8 | 0 | 33 |
| 17.3 | 8.2 | FxM | 16 | 11 | 13 | 1 | 41 |
| 17.6 | 8.3 | FxM | 8 | 11 | 24 | 0 | 43 |
| 2.7 | 24.1 | FxM | 27 | 17 | 4 | 1 | 49 |
| 26.4 | 8.3 | FxM | 17 | 7 | 11 | 1 | 36 |
| 27.6 | 6.1 | FxH | 24 | 27 | 5 | 0 | 56 |
| 4.1 | 8.3 | FxM | 14 | 8 | 13 | 0 | 35 |
| 10.1x8.3 | 27.6x6.1 | FxM | 26 | 5 | 19 | 6 | 56 |
| 14.1x6.1 | 16.1x24.1 | FxM | 30 | 16 | 46 | 2 | 94 |
| 15.5x6.1 | 16.1x24.1 | FxM | 34 | 7 | 30 | 3 | 74 |
| 16.1x24.1 | 26.4x8.3 | FxM | 8 | 8 | 11 | 2 | 29 |
| 2.7x24.1 | 2.7x24.1 | HxH | 13 | 34 | 0 | 0 | 47 |
| 27.6x6.1 | 17.6x8.3 | FxM | 53 | 15 | 40 | 26 | 134 |
| 27.6x6.1 | 4.1x8.3 | FxH | 14 | 24 | 0 | 0 | 38 |
| 13.2 | 6.1 | FxH | 5 | 23 | 0 | 0 | 28 |
| 17.6 | 8.3 | FxM | 17 | 5 | 17 | 0 | 39 |
| 10.1 | 8.3 | FxM | 13 | 6 | 13 | 0 | 32 |
| 14.1 | 6.1 | FxH | 13 | 20 | 0 | 0 | 33 |
| 14.1 | 6.1 | FxH | 38 | 98 | 0 | 0 | 136 |
| 10.1 | 8.3 | FxM | 5 | 10 | 7 | 0 | 22 |
| 4.1 | 8.3 | FxM | 9 | 8 | 13 | 0 | 30 |
| 25.1 | 25.1 | HxH | 12 | 42 | 0 | 0 | 54 |
| 23.11 | 25.1 | FxH | 19 | 31 | 0 | 0 | 50 |
| 17.5 | 25.1 | FxH | 11 | 19 | 0 | 0 | 30 |
| 12.16 | 12.16 | HxH | 5 | 17 | 0 | 0 | 22 |
| 4.3 | 12.16 | FxH | 25 | 56 | 0 | 0 | 81 |
| 22.15 | 12.16 | FxH | 32 | 71 | 0 | 0 | 103 |
| 18.3 | 18.3 | HxH | 23 | 69 | 0 | 0 | 92 |
| 3.17 | 18.3 | FxH | 21 | 47 | 0 | 0 | 68 |
| 11.2 | 18.3 | FxH | 17 | 38 | 0 | 0 | 55 |
| 26.8 | 18.3 | FxH | 13 | 29 | 0 | 0 | 42 |
| 12.1 | 4.5 | FxM | 17 | 10 | 30 | 3 | 60 |
| 7.1 | 4.5 | FxM | 32 | 18 | 45 | 2 | 97 |
| 18.9 | 11.4 | FxM | 12 | 8 | 18 | 0 | 38 |
| 22.3 | 11.4 | FxM | 9 | 5 | 10 | 0 | 24 |
| 17.6 | 11.4 | FxM | 18 | 11 | 23 | 0 | 52 |
| 9.1 | 25.1 | FxM | 26 | 16 | 19 | 0 | 61 |
| 12.4 | 25.1 | FxM | 29 | 14 | 20 | 1 | 64 |
| 23.12 | 25.1 | FxM | 16 | 10 | 20 | 0 | 46 |
| 24.3 | 19.3 | FxM | 12 | 5 | 9 | 0 | 26 |
| 11.2 | 19.3 | FxM | 19 | 9 | 30 | 4 | 62 |
| 15.3 | 19.3 | FxM | 25 | 12 | 33 | 2 | 72 |
| 132.10.1 | 132.24.1 | FxM | 20 | 4 | 26 | 1 | 51 |
| 132.20.1 | 132.8.3 | FxM | 13 | 3 | 14 | 0 | 30 |
| 132.12.5 | 132.8.3 | FxM | 21 | 13 | 29 | 0 | 63 |
| 132.28.1 | 132.8.1 | FxM | 19 | 8 | 23 | 0 | 50 |
| 132.7.8 | 132.8.1 | FxM | 12 | 4 | 20 | 0 | 36 |
| 132.14.8 | 132.8.1 | FxM | 6 | 5 | 14 | 0 | 25 |
| 132.5.17 | 132.17.1 | FxM | 18 | 7 | 23 | 1 | 49 |
| 132.18.3 | 132.17.1 | FxM | 23 | 10 | 30 | 1 | 64 |
| 132.25.9 | 132.17.1 | FxM | 15 | 4 | 20 | 0 | 39 |
| 132.5.3 | 132.5.3 | HxH | 17 | 50 | 0 | 0 | 67 |
| 132.14.1 | 132.5.3 | FxH | 12 | 20 | 0 | 0 | 32 |
| 132.11.8 | 132.5.3 | FxH | 9 | 13 | 0 | 0 | 22 |
| 132.25.3 | 132.5.3 | FxH | 21 | 37 | 0 | 0 | 58 |
| 132.19.1 | 132.19.1 | HxH | 12 | 30 | 0 | 0 | 42 |
| 132.7.4 | 132.19.1 | FxH | 15 | 28 | 0 | 0 | 43 |
| 132.21.2 | 132.19.1 | FxH | 12 | 29 | 0 | 0 | 41 |

**Table S4.** The number of males, sterile males, females and hermaphrodites in the offspring of 39 FxM crosses used for recombination rate fitting, and the *p-values* of fitting each color-coded scenario (see figure 5) to the sex ratio of each cross.

| Cross | Mother | Father | F | H | M | N | Total | Blue | Red | Green | Black | Orange | Pink |
| --- | --- | --- | --- | --- | --- | --- | --- | --- | --- | --- | --- | --- | --- |
| 1 | 10.1 | 24.1 | 10 | 6 | 9 | 1 | 26 | 0.272 | 0 | 0.444 | 0.005 | 0 | 0 |
| 2 | 10.1 | 8.3 | 13 | 6 | 7 | 0 | 26 | 0.021 | 0 | 0.084 | 0 | 0 | 0 |
| 3 | 13.2 | 8.3 | 22 | 11 | 13 | 5 | 51 | 0.158 | 0 | 0 | 0 | 0 | 0 |
| 4 | 16.1 | 24.1 | 15 | 3 | 10 | 1 | 29 | 0.222 | 0 | 0.126 | 0 | 0 | 0 |
| 5 | 16.1 | 8.2 | 13 | 8 | 14 | 4 | 39 | 0.612 | 0 | 0.011 | 0 | 0 | 0 |
| 6 | 17.1 | 8.2 | 3 | 22 | 8 | 0 | 33 | 0 | 0 | 0 | 0.006 | 0.003 | 0 |
| 7 | 17.3 | 8.2 | 16 | 11 | 13 | 1 | 41 | 0.02 | 0 | 0.136 | 0 | 0 | 0 |
| 8 | 17.6 | 8.3 | 8 | 11 | 24 | 0 | 43 | 0 | 0 | 0.081 | 0.049 | 0 | 0 |
| 9 | 26.4 | 8.3 | 17 | 7 | 11 | 1 | 36 | 0.091 | 0 | 0.147 | 0 | 0 | 0 |
| 10 | 4.1 | 8.3 | 14 | 8 | 13 | 0 | 35 | 0.015 | 0 | 0.375 | 0 | 0 | 0 |
| 11 | 10.1x8.3 | 27.6x6.1 | 26 | 5 | 19 | 6 | 56 | 0.493 | 0 | 0 | 0 | 0 | 0 |
| 12 | 14.1x6.1 | 16.1x24.1 | 30 | 16 | 46 | 2 | 94 | 0 | 0 | 0.964 | 0 | 0 | 0 |
| 13 | 15.5x6.1 | 16.1x24.1 | 34 | 7 | 30 | 3 | 74 | 0.035 | 0 | 0.018 | 0 | 0 | 0 |
| 14 | 16.1x24.1 | 26.4x8.3 | 8 | 8 | 11 | 2 | 29 | 0.161 | 0 | 0.119 | 0.099 | 0 | 0 |
| 15 | 27.6x6.1 | 17.6x8.3 | 53 | 15 | 40 | 26 | 134 | 0.103 | 0 | 0 | 0 | 0 | 0 |
| 16 | 17.6 | 8.3 | 17 | 5 | 17 | 0 | 39 | 0.017 | 0 | 0.4 | 0 | 0 | 0 |
| 17 | 10.1 | 8.3 | 13 | 6 | 13 | 0 | 32 | 0.04 | 0 | 0.624 | 0 | 0 | 0 |
| 18 | 10.1 | 8.3 | 5 | 10 | 7 | 0 | 22 | 0 | 0 | 0.017 | 0.196 | 0.002 | 0 |
| 19 | 4.1 | 8.3 | 9 | 8 | 13 | 0 | 30 | 0.011 | 0 | 0.475 | 0.025 | 0 | 0 |
| 20 | 12.1 | 4.5 | 17 | 10 | 30 | 3 | 60 | 0.045 | 0 | 0.324 | 0 | 0 | 0 |
| 21 | 7.1 | 4.5 | 32 | 18 | 45 | 2 | 97 | 0 | 0 | 0.922 | 0 | 0 | 0 |
| 22 | 18.9 | 11.4 | 12 | 8 | 18 | 0 | 38 | 0.007 | 0 | 0.783 | 0.001 | 0 | 0 |
| 23 | 22.3 | 11.4 | 9 | 5 | 10 | 0 | 24 | 0.093 | 0 | 0.78 | 0.004 | 0 | 0 |
| 24 | 17.6 | 11.4 | 18 | 11 | 23 | 0 | 52 | 0.001 | 0 | 0.584 | 0 | 0 | 0 |
| 25 | 9.1 | 25.1 | 26 | 16 | 19 | 0 | 61 | 0 | 0 | 0.015 | 0 | 0 | 0 |
| 26 | 12.4 | 25.1 | 29 | 14 | 20 | 1 | 64 | 0.002 | 0 | 0.042 | 0 | 0 | 0 |
| 27 | 23.12 | 25.1 | 16 | 10 | 20 | 0 | 46 | 0.003 | 0 | 0.586 | 0 | 0 | 0 |
| 28 | 24.3 | 19.3 | 12 | 5 | 9 | 0 | 26 | 0.066 | 0 | 0.34 | 0 | 0 | 0 |
| 29 | 11.2 | 19.3 | 19 | 9 | 30 | 4 | 62 | 0.158 | 0 | 0.1 | 0 | 0 | 0 |
| 30 | 15.3 | 19.3 | 25 | 12 | 33 | 2 | 72 | 0.014 | 0 | 0.856 | 0 | 0 | 0 |
| 31 | 132.10.1 | 132.24.1 | 20 | 4 | 26 | 1 | 51 | 0.012 | 0 | 0.271 | 0 | 0 | 0 |
| 32 | 132.20.1 | 132.8.3 | 13 | 3 | 14 | 0 | 30 | 0.043 | 0 | 0.413 | 0 | 0 | 0 |
| 33 | 132.12.5 | 132.8.3 | 21 | 13 | 29 | 0 | 63 | 0 | 0 | 0.582 | 0 | 0 | 0 |
| 34 | 132.28.1 | 132.8.1 | 19 | 8 | 23 | 0 | 50 | 0.004 | 0 | 0.674 | 0 | 0 | 0 |
| 35 | 132.7.8 | 132.8.1 | 12 | 4 | 20 | 0 | 36 | 0.008 | 0 | 0.561 | 0 | 0 | 0 |
| 36 | 132.14.8 | 132.8.1 | 6 | 5 | 14 | 0 | 25 | 0.018 | 0 | 0.628 | 0.061 | 0 | 0 |
| 37 | 132.5.17 | 132.17.1 | 18 | 7 | 23 | 1 | 49 | 0.045 | 0 | 0.928 | 0 | 0 | 0 |
| 38 | 132.18.3 | 132.17.1 | 23 | 10 | 30 | 1 | 64 | 0.007 | 0 | 0.954 | 0 | 0 | 0 |
| 39 | 132.25.9 | 132.17.1 | 15 | 4 | 20 | 0 | 39 | 0.009 | 0 | 0.451 | 0 | 0 | 0 |

